# Loss of CFIm activates YAP/TAZ and connects mRNA cleavage and polyadenylation inhibition to BRCAness

**DOI:** 10.1101/2025.10.21.683728

**Authors:** Madeleine Goldthorpe, Jiahui Hou, Abhi Nadendla, Hatice Ulku Osmanbeyoglu, Hun-Way Hwang

## Abstract

The YAP/TAZ transcription program plays a crucial role in development, regeneration, and cancer. CFIm is a master regulator of mRNA alternative polyadenylation (APA). Loss of CFIm function was shown in human cancer but its impact on the gene expression program is not well characterized. Here we report the discovery that loss of CFIm in cancer cells activates YAP/TAZ and promotes therapeutic resistance. We identified a CFIm-NEDD4L-LATS1 regulatory axis that suppressed YAP/TAZ activation. Furthermore, we found that CFIm loss in the presence of mRNA cleavage and polyadenylation (CPA) inhibition repressed key DNA repair genes in the Fanconi anemia (FA) and homology-directed repair (HDR) pathways, which induced the BRCAness phenotype and aggravated DNA damage. Our study reveals a hidden link between mRNA 3′ end processing and the YAP/TAZ transcription program in addition to illustrating how CFIm function impacts the selection of therapeutic agents in cancer.

## INTRODUCTION

Yes-associated protein (YAP) and transcriptional co-activator with PDZ-binding motif (TAZ) are two homologous transcriptional co-activators that do not directly bind to DNA but are crucial for normal development and tissue regeneration^1^. YAP/TAZ activation is seen in a wide range of human cancer, and activation of YAP/TAZ and their transcriptional program in cancer is an important mechanism for the development of therapeutic resistance and disease progression^2^. Therefore, it is essential to develop a thorough understanding in how cancer cells activate YAP/TAZ. YAP/TAZ is normally repressed by the mammalian Hippo pathway, which is a cascade consisting of two kinases, MST1/2 and LATS1/2, in addition to multiple adaptor proteins^3^. LATS1/2 directly phosphorylates YAP/TAZ to suppress their nuclear translocation and promote their degradation^1^. However, with the exception of certain cancer types such as malignant mesothelioma, meningioma and Schwannoma, mutations in the Hippo pathway components are not common in human cancer^4^. This suggests that cancer cells have alternative ways to activate YAP/TAZ to their advantages without shutting down the Hippo pathway. For example, more than 40% of uveal melanoma carry activating mutations in G-protein subunits *GNAQ* or *GNA11*^5,6^, and it was found that these mutations drive YAP/TAZ activation to promote tumorigenesis^7^.

In humans, the mRNA 3′-end processing complex consists of more than 20 proteins that are grouped into 4 factors: CPSF, CSTF, CFIIm and CFIm^8^. The human CFIm consists of two subunits^9^: a small subunit, CFIm25 (encoded by the *NUDT21* gene), which directly binds to the UGUA motif in the polyadenylation site (PAS)^9^, and two alternative large subunits, CFIm59 (encoded by the *CPSF7* gene) and CFIm68 (encoded by the *CPSF6* gene), which activate the mRNA 3′ end processing by interacting with CPSF^10^. However, CFIm68 is a much stronger activator of the mRNA 3′-end processing than CFIm59^10^. CFIm is not essential for the cleavage reaction in mRNA 3′ end processing^11^, but it has a strong influence on PAS choices. We and others reported predominantly (>90%) proximal APA shifts in hundreds of genes from loss of either NUDT21 or CPSF6^10,12–14^. However, how these widespread APA changes from CFIm loss regulate the activities of cell signaling pathways is not fully understood, but the impact of CFIm on cell signaling starts to emerge from recent studies. For example, in HEK293 cells, CFIm promoted the activities of ERK1/2, which were correlated with the expression level of CFIm^12^. In contrast, in human primary fibroblasts, NUDT21 knockdown activated fibrotic pathways that promoted the expression of profibrotic genes^15^. CPSF3 is the endonuclease in the mRNA 3′-end processing complex^16^. CPSF3 inhibition suppresses mRNA CPA, which results in transcription readthrough and APA changes^17,18^. The first CPSF3 inhibitor, JTE-607, was recently shown to have anti-proliferation activities in some types of cancer such as acute myeloid leukemia (AML) and Ewing’s sarcoma^19^.

The tumor suppressors BRCA1 and BRCA2 both play key roles in the homology-directed repair (HDR) pathways to maintain genome stability^20^. *BRCA1/2* mutant cancer is deficient in HDR, and it has increased vulnerability to poly(adenosine 5′-diphosphate–ribose) polymerase (PARP) inhibitors and DNA crosslinking agents such as platinum salts and mitomycin C (MMC)^21^. The term “BRCAness” was used to describe the similar phenotype (sensitivity to PARP inhibitors) observed in cancer without *BRCA1/2* germline mutations^22^. The presence of HDR defects is now considered as the mechanism for the BRCAness phenotype, and additional genes that cause BRCAness when mutated in cancer have been identified^20,21^.

CFIm loss-of-function (LOF), either caused by low expression or inhibitory phosphorylation, has been reported in several types of cancer^23–25^. In this study, we used two different cancer cell models to investigate how CFIm loss affects the cell signaling activities. We discovered that CFIm loss activated YAP/TAZ and their target genes, and we identified LATS1 and NEDD4L as the key genes linking CFIm loss to YAP/TAZ activation. In search for a therapeutic strategy against CFIm loss, we found that CFIm loss increased sensitivity to mRNA CPA inhibition. Furthermore, we demonstrated that the combination of CFIm loss and CPSF3 inhibition induced BRCAness. Overall, our study identified restraining the YAP/TAZ transcriptional program and supporting DNA damage repair as previously unknown tumor suppressive roles of CFIm.

## RESULTS

### CFIm loss activates YAP/TAZ in colon and breast cancer cells

To study the biological function of CFIm (Fig. 1A), we used the degradation tag (dTAG) system^26^, which can achieve a rapid, inducible and specific depletion of the target protein. In the presence of a chemical inducer such as dTAG^V^-1, the FKBP12^F36V^ degron-tagged target protein binds to an endogenous E3 ubiqutin ligase and is subsequently degraded^26^. We chose the HCT116 colon cancer cells as the model because of its favorable characteristics: a near-diploid genome that is amenable to genome editing with the clustered regularly interspaced short palindromic repeats (CRISPR) technology^27^ and the reported success in implementing the dTAG system^28^. We generated a single-cell clone of HCT116 colon cancer cell in which both alleles of *NUDT21* were tagged with the FKBP12^F36V^ degron tag by CRISPR-Cas9 editing (referred to as dTAG-NUDT21 HCT116 cells hereafter) (Fig. S1A). The tagging of *NUDT21* did not substantially change NUDT21 and CPSF6 protein expression when compared with wildtype HCT116 cells (Fig. S1B). Treating dTAG-NUDT21 HCT116 cells with dTAG^V^-1 resulted in complete depletion of NUDT21 proteins within 4 hours (Fig. S1C). We also verified that dTAG^V^-1 treatment in dTAG-NUDT21 HCT116 cells resulted in proximal APA shift of *TIMP2*, a known CFIm target^13,29^ (Fig. S1D).

**Figure 1.**
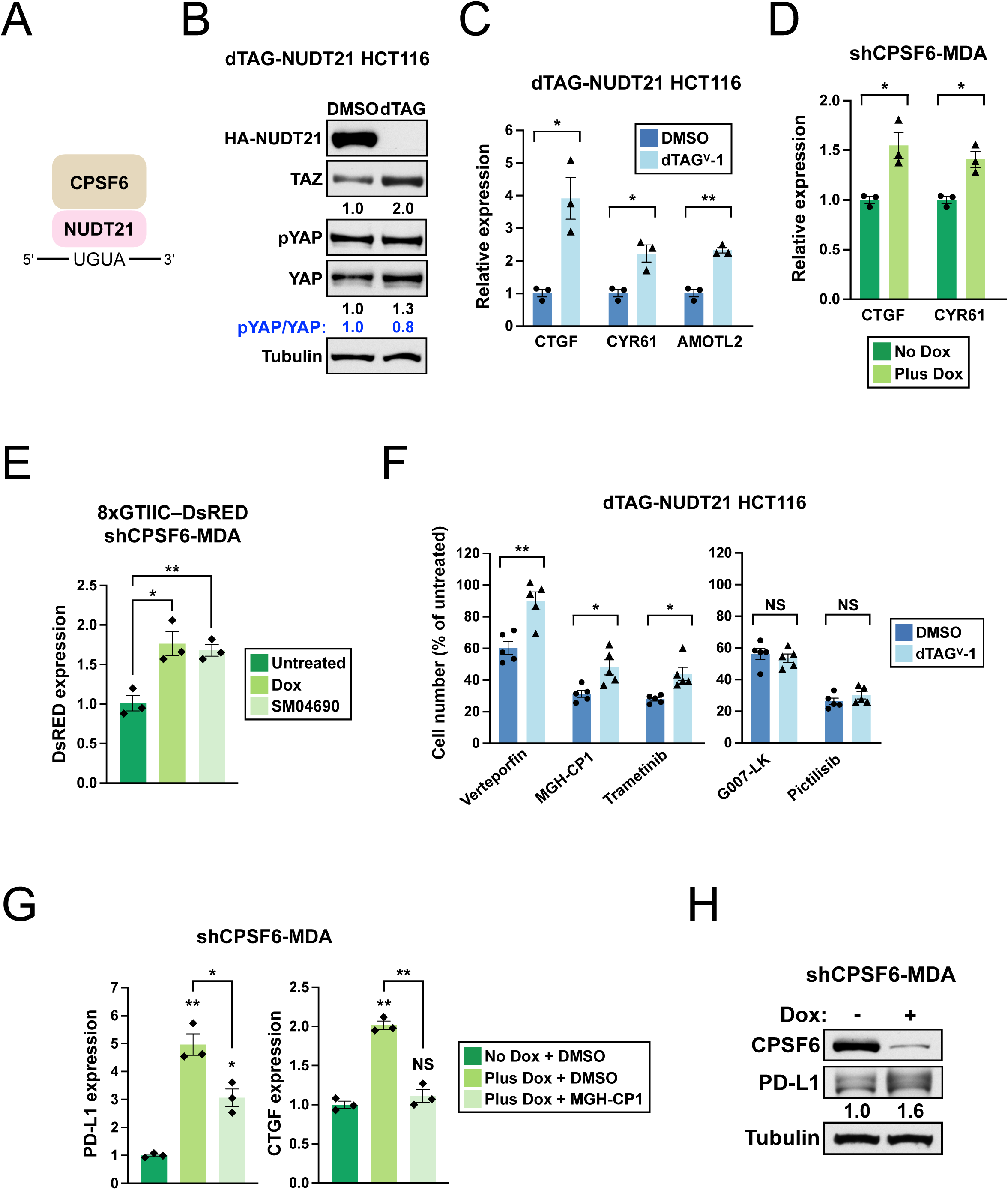
Loss of CFIm activates YAP/TAZ signaling and promotes therapeutic resistance. (**A**) Illustrations showing the human CFIm. UGUA: CFIm binding motif. (**B**) Western blots showing higher YAP/TAZ expression and a lower Phospho-YAP (Ser127) (pYAP)/YAP ratio in dTAG-NUDT21 HCT116 cells treated with dTAG^V^-1 for 4 h. Tubulin: loading control. (**C**) Bar graphs showing the expression of YAP/TAZ targets measured by RT-qPCR in dTAG-NUDT21 HCT116 cells treated with either DMSO or dTAG^V^-1 for 24 h from 3 independent experiments (n=3). (**D**) Bar graphs showing the expression of YAP/TAZ targets measured by RT-qPCR in shCPSF6-MDA cells treated with doxycycline (Plus Dox) or remained untreated (No Dox) for 72 h from 3 independent experiments (n=3). (**E**) Bar graphs showing DsRED expression measured by RT-qPCR in 8xGTIIC–DsRED shCPSF6-MDA cells receiving different treatments for 72 h from 3 independent experiments (n=3). (**F**) Bar graphs showing the relative number of live dTAG-NUDT21 HCT116 cells treated with different reagents in combination with either DMSO or dTAG^V^-1 for 48 h from 5 independent experiments (n=5). (**G**) Bar graphs showing PD-L1 (left) or CTGF (right) expression measured by RT-qPCR in shCPSF6-MDA cells receiving different treatments for 72 h from 3 independent experiments (n=3). (**H**) Western blots showing higher PD-L1 expression in shCPSF6-MDA cells with doxycycline treatment for 72 h. dTAG^V^-1: 500nM. Doxycycline: 1 μg/mL. Verteporfin: 5μM. MGH-CP1: 10μM. Trametinib: 10nM. G007-LK: 10μM. Pictilisib: 1μM. SM04690: 100nM. Error bars indicate SEM. Statistical significance is determined by two-tailed t-test. NS: not significant, *: p<0.05, **: p<0.01.

Next, we treated dTAG-NUDT21 HCT116 cells with either DMSO or dTAG^V^-1 to investigate how CFIm loss impacts major cell signaling pathways. We found that dTAG^V^-1 treatment increased YAP and TAZ protein expression without any changes in phospho-YAP (inactive form) abundance, suggesting possible YAP/TAZ activation from CFIm loss (Fig. 1B). We also performed RT-qPCR experiments in dTAG^V^-1 treated dTAG-NUDT21 HCT116 cells to examine the expression of YAP/TAZ direct target genes–*CTGF*, *CYR61* and *AMOTL2*^30^. dTAG^V^-1 treatment increased the expression of all 3 genes, indicating that the YAP/TAZ transcription program was likely activated in response to CFIm loss (Fig. 1C). To examine the relationship between CFIm loss and YAP/TAZ in a distinct biological context with a different gene inhibition technique, we next used an MDA-MB-231 breast cancer cell line that was previously generated in our lab and carries a doxycycline-inducible short hairpin RNA (shRNA) targeting CPSF6^31^ (referred to as shCPSF6-MDA cells hereafter). Activation of the CPSF6 shRNA by doxycycline treatment for 72 hours in shCPSF6-MDA cells strongly reduced CPSF6 expression but did not result in its complete loss (Fig. S1E). Next, we grew shCPSF6-MDA cells in the presence or absence of doxycycline for 72 hours, and we performed RT-qPCR to examine the expression of YAP/TAZ target genes. Consistent with the results in dTAG-NUDT21 HCT116 cells, we found that the expression of *CTGF* and *CYR61* was again increased in shCPSF6-MDA cells with low CPSF6 expression (Fig. 1D). Lastly, we stably transduced shCPSF6-MDA cells with a YAP/TAZ reporter (8xGTIIC–DsRED)^32^ to examine whether the upregulation of YAP/TAZ targets occurs at the transcription level. The induction of DsRED expression was similar between doxycycline and SM04690, a YAP activator^33^ (Fig. 1E), confirming the transcriptional activation effects on a YAP/TAZ-responsive promoter from CPSF6 knockdown. Taken together, our results show that CFIm loss activates YAP/TAZ in colon and breast cancer cells.

### CFIm loss promotes therapeutic resistance and PD-L1 expression

YAP/TAZ activation in multiple types of cancer cells increased resistance to different therapeutic agents^2,4^. Therefore, we next performed cell proliferation assay to examine whether CFIm loss changes the sensitivity of dTAG-NUDT21 HCT116 cells to different classes of therapeutic agents (verteporfin and MGH-CP1: YAP/TAZ inhibitors, Trametinib: MEK inhibitor; G007-LK: WNT pathway inhibitor; pictilisib: PI3K inhibitor). We found that dTAG^V^-1-treated dTAG-NUDT21 HCT116 cells were more resistant to Trametinib (Fig. 1F). This finding is consistent with a previous report that YAP promotes resistance to MEK inhibitors^34^. In contrast, the toxicity of G007-LK and pictilisib were similar between the DMSO and dTAG^V^-1 groups. Unexpectedly, dTAG^V^-1-treated dTAG-NUDT21 HCT116 cells were also more resistant to both YAP/TAZ inhibitors, verteporfin and MGH-CP1 (Fig. 1F), suggesting the possibility of additional unidentified signaling changes that promote resistance to YAP/TAZ inhibitors^35,36^.

PD-L1 is an immune checkpoint protein that cancer cells use to suppress T-cell activity and evade immunotherapy^37^. In breast cancer cells, TAZ activation increases PD-L1 expression and suppresses T-cell function^38^. Therefore, we next examined PD-L1 expression in shCPSF6-MDA cells grown in the presence or absence of doxycycline. Consistent with the previous report, we found increased expression of *PD-L1* mRNA (Fig. 1G) and protein (Fig. 1H) in doxycycline-treated shCPSF6-MDA cells. Interestingly, treatment with MGH-CP1 completely abolished the increase in *CTGF* and *CYR61* mRNA expression from doxycycline-induced CPSF6 knockdown (Fig. 1G, right panel, and Fig. S1F) but it only partially rescued *PD-L1* mRNA expression (Fig. 1G, left panel). These results indicate that the increase in *CTGF* and *CYR61* expression from CFIm loss is solely caused by higher YAP/TAZ transcriptional activities while the increase in *PD-L1* expression has other contributory mechanisms. For example, post-transcriptional regulation of *PD-L1* expression through 3′ UTR was previously reported^37,39^. Altogether, we demonstrate that cancer cells that lose CFIm become more resistant to different classes of chemical inhibitors and have stronger *PD-L1* expression, which is beneficial for evading immune responses.

### LATS1/2 are required for activation of YAP/TAZ from CFIm loss

We next sought to investigate the mechanistic link between CFIm loss and YAP/TAZ activation. We first compiled a list of established YAP/TAZ regulators to examine their possible involvement in YAP/TAZ activation from CFIm loss in dTAG-NUDT21 HCT116 cells (Fig. 2A). We used siRNAs or chemical inhibitors to inhibit the function of each YAP/TAZ regulators and performed RT-qPCR to assay YAP/TAZ activation using *CTGF* mRNA expression as a surrogate. Because YAP/TAZ activities are regulated by cell density^40^, we first examined whether cell density affects YAP/TAZ activation from CFIm loss. We found that CFIm loss activated YAP/TAZ in both confluent and sub-confluent cells, indicating that YAP/TAZ activation from CFIm loss is independent of cell density (Fig. 2B). Next, we examined the roles of MST1/2 and NF2, both of which are YAP repressors in the Hippo pathway^1^. Treatment with XMU-MP-1, an MST1/2 inhibitor^41^, and siRNA-mediated knockdown of NF2 (Fig. S2A) both elevated the baseline CTGF expression, as was expected from the inhibition of a YAP repressor (Fig. 2C). In contrast, treatment with a CK2 inhibitor, CX4945, lowered the baseline CTGF expression as was expected from the inhibition of CK2, a YAP activator outside of the Hippo pathway^42^ (Fig. 2D). However, none of the 3 treatments blocked the increase in CTGF expression, indicating that MST1/2, NF2, and CK2 are not required for YAP/TAZ activation from CFIm loss.

**Figure 2.**
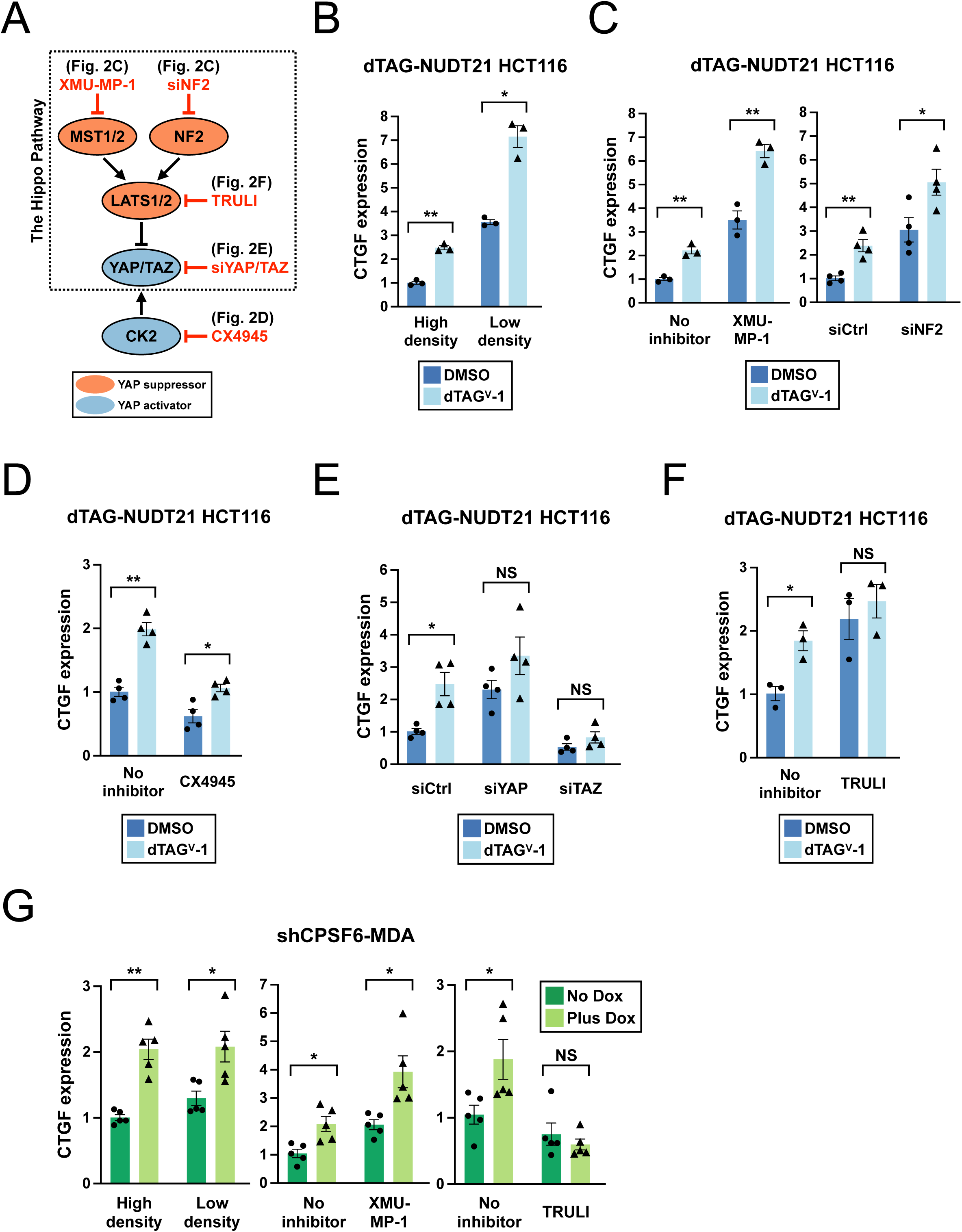
LATS1/2 are required for activation of YAP/TAZ signaling from CFIm loss. (**A**) An overview of the YAP/TAZ regulators, the chemical inhibitors or siRNAs used for inhibition, and the figure panels showing the experimental results. Orange: YAP signaling suppressors. Blue: YAP signaling activators. (**B**) Bar graphs showing CTGF expression measured by RT-qPCR in dTAG-NUDT21 HCT116 cells plated at different densities followed by DMSO or dTAG^V^-1 treatment for 24 h from 3 independent experiments (n=3). (**C**) Bar graphs showing CTGF expression measured by RT-qPCR in dTAG-NUDT21 HCT116 cells. (Left panel) Cells were treated with DMSO alone, dTAG^V^-1 alone, DMSO plus XMU-MP-1 or dTAG^V^-1 plus XMU-MP-1 for 24 h from 3 independent experiments (n=3). (Right panel) Cells were first transfected with control siRNAs (siCtrl) or siRNAs targeting NF2 (siNF2) for 48 h and then were treated with DMSO or dTAG^V^-1 for 24 h before RNA collection from 4 independent experiments (n=4). (**D**∼**F**) Bar graphs showing CTGF expression measured by RT-qPCR in dTAG-NUDT21 HCT116 cells. (**D**) Cells were treated with DMSO alone, dTAG^V^-1 alone, DMSO plus CX4945 or dTAG^V^-1 plus CX4945 for 24 h from 4 independent experiments (n=4). (**E**) Cells were first transfected with control siRNAs (siCtrl), siRNAs targeting YAP (siYAP), or siRNAs targeting TAZ (siTAZ) for 48 h and then were treated with DMSO or dTAG^V^-1 for 24 h before RNA collection from 4 independent experiments (n=4). (**F**) Cells were treated with DMSO alone, dTAG^V^-1 alone, DMSO plus TRULI or dTAG^V^-1 plus TRULI for 24 h from 4 independent experiments (n=4). (**G**) Bar graphs showing CTGF expression measured by RT-qPCR in shCPSF6-MDA cells. (Left panel) Cells were plated at different densities followed by doxycycline treatment (Plus Dox) or remained untreated (No Dox) for 72 h from 4 independent experiments (n=4). (Middle panel) Cells were untreated, or treated with doxycycline alone, XMU-MP-1 alone, and doxycycline plus XMU-MP-1 for 72 h from 5 independent experiments (n=5). (Right panel) Cells were untreated, or treated with doxycycline alone, TRULI alone, and doxycycline plus TRULI for 72 h from 5 independent experiments (n=5). In (B) and (G), cells plated at high density reached complete confluence and cells plated at low density remained <50% confluence at RNA collection. XMU-MP-1: 2μM. CX4945: 2μM. TRULI: 10μM. Error bars indicate SEM. Statistical significance is determined by two-tailed t-test. NS: not significant, *: p<0.05, **: p<0.01.

Next, we separately inhibited YAP and TAZ expression using siRNAs (Fig. S2B). Notably, the knockdown efficiency was lower for YAP compared with TAZ, which was likely due to a stronger role of YAP in HCT116 cell proliferation^43^. Nevertheless, both YAP and TAZ knockdown suppressed the increase in CTGF expression (Fig. 2E), supporting an essential role for both YAP and TAZ as expected. Lastly, we examined the role of LATS1/2 using TRULI, a specific LATS1/2 inhibitor^44^. TRULI increased the baseline CTGF expression as expected from inhibition of LATS1/2 (Fig. 2F). Furthermore, TRULI very strongly suppressed the increase in CTGF expression (Fig. 2F), indicating that LATS1/2 are necessary for YAP/TAZ activation from CFIm loss.

To ensure these results are not specific to dTAG-NUDT21 HCT116 cells, we also examined the roles of cellular density, MST1/2, and LATS1/2 in YAP/TAZ activation from CFIm loss in shCPSF6-MDA cells. As in dTAG-NUDT21 HCT116 cells, the increase in CTGF expression from CFIm loss requires LATS1/2 but it is independent of cell density and MST1/2 in shCPSF6-MDA cells (Fig. 2G). Taken together, our results from two different cell models indicated that LATS1/2 are required for YAP/TAZ activation from CFIm loss.

### TAZ mRNA 3′ UTR shortening from CFIm loss increases its expression

In human, both YAP (encoded by the *YAP1* gene) and TAZ (encoded by the *WWTR1* gene) have short 3′ UTR mRNA isoforms (Fig. 3A). 3′ UTR shortening has been shown to relieve repressive regulation and increase gene expression^45^. To further explore the mechanistic link between CFIm loss and YAP/TAZ activation, we next investigated the possibility that APA changes from CFIm loss contribute to higher expression of YAP and TAZ (Fig. 1B). We performed RT-qPCR experiments in both dTAG-NUDT21 HCT116 and shCPSF6-MDA cells to examine whether CFIm loss results in 3′ UTR shortening in YAP and TAZ. Indeed, CFIm loss resulted in a 90% or higher decrease in full-length (FL) 3′ UTR mRNA isoform for YAP and TAZ in both cell lines (Fig. 3B-C). We next adopted a previously published dual fluorescence reporter^46^ to measure how 3′ UTR shortening affects YAP and TAZ expression by flow cytometry (Fig. 3D). In this assay, we first cloned the 3′ UTR of interest into the empty reporter downstream of the mCherry coding sequence. We next transiently expressed the resulting reporter in HEK293T cells to measure the mCherry/GFP ratio, which reported the effects from the 3′ UTR of interest on gene expression. For YAP, the FL 3′ UTR reporter and the short 3′ UTR reporter generated similar mCherry/GFP ratios, suggesting that the 3′ UTR shortening from CFIm loss is less likely to alter YAP expression (Fig. 3E). In contrast, for TAZ, the FL 3′ UTR lowered mCherry expression compared with the short 3′ UTR (mCherry/GFP: FL, 0.71; short, 0.92) (Fig. 3E). In summary, we discovered that CFIm promotes the usage of full-length 3′ UTR in YAP and TAZ, and we further demonstrated that TAZ mRNA 3′ UTR shortening contributes to its increased expression with CFIm loss.

**Figure 3.**
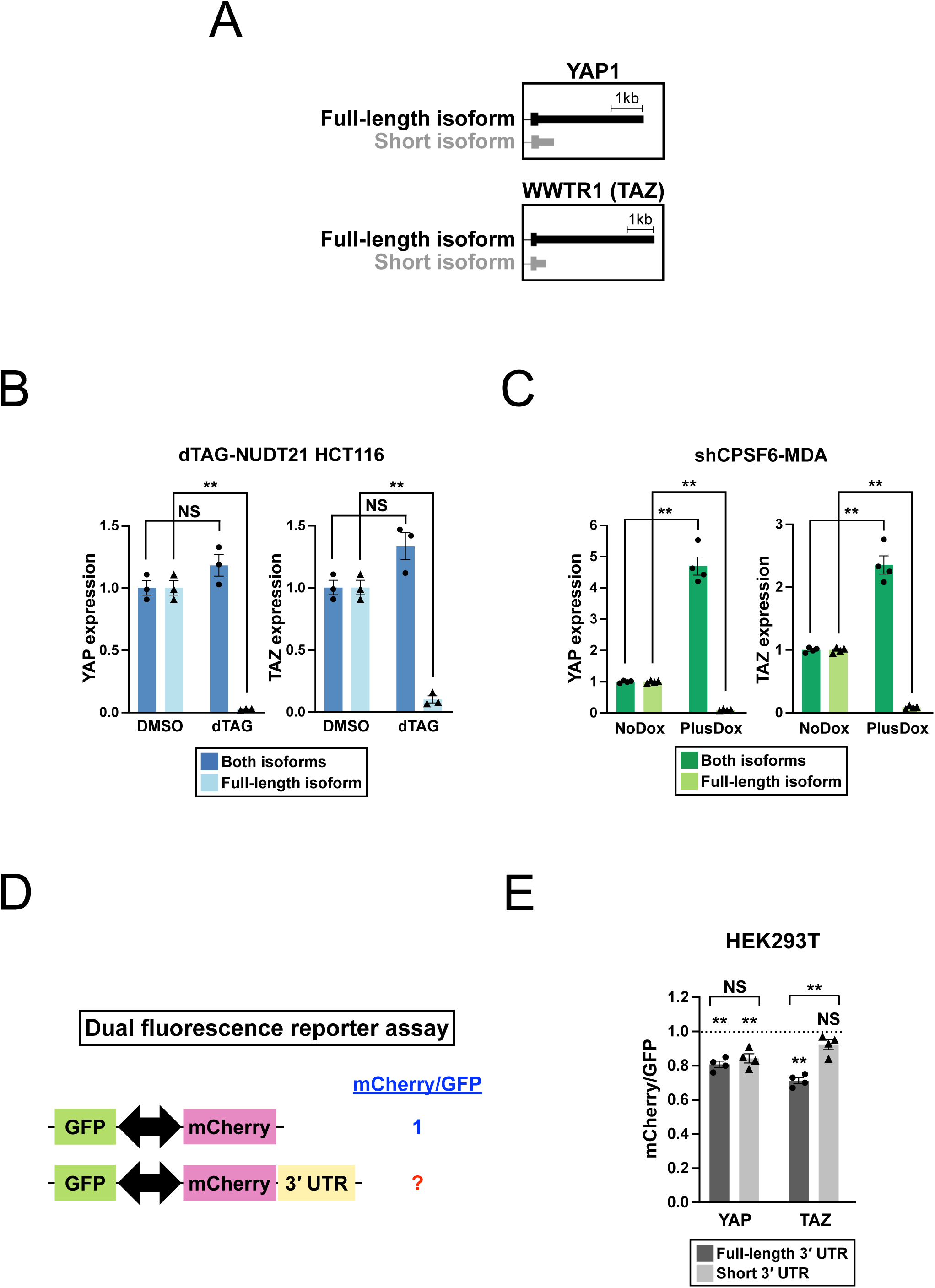
TAZ mRNA 3′ UTR shortening from CFIm loss contributes to its increased expression. (**A**) Illustrations showing the full-length 3′ UTR and short 3′ UTR mRNA isoforms of YAP and TAZ as supported by Ensembl annotations and PAPERCLIP experiments. (**B**) Bar graphs showing expression of YAP and TAZ mRNA isoforms measured by RT-qPCR in dTAG-NUDT21 HCT116 cells treated with DMSO or dTAG^V^-1 for 24 h from 3 independent experiments (n=3). (**C**) Bar graphs showing expression of YAP and TAZ mRNA isoforms measured by RT-qPCR in shCPSF6-MDA cells treated with doxycycline (Plus Dox) or remained untreated (No Dox) for 72 h from 4 independent experiments (n=4). (**D**) Illustration showing the dual fluorescence reporter assay used to measure the effects from the 3′ UTR of interest on gene expression by flow cytometry. The mCherry/GFP ratio from the empty vector is set to 1. Arrows indicate the bi-directional promoter. (**E**) Bar graphs showing the mCherry/GFP ratio from YAP and TAZ 3′ UTR reporters in 293T cells from 4 independent experiments (n=4). Error bars indicate SEM. Statistical significance is determined by two-tailed t-test. NS: not significant, **: p<0.01.

### Identification of a CFIm-NEDD4L-LATS1 regulatory axis

The E3 ubiquitin ligases NEDD4 and NEDD4L were previously reported to activate YAP in different biological contexts^47–49^. Therefore, we next explored the possibility that NEDD4 and/or NEDD4L participated in YAP/TAZ activation from CFIm loss. We first examined our previously published PAPERCLIP profiling datasets^13,31^ to see whether CFIm regulates the APA of NEDD4 and/or NEDD4L. We found that CPSF6 knockdown resulted in strong 3′ UTR shortening in both NEDD4 and NEDD4L (Fig. 4A). We next performed RT-qPCR experiments in dTAG-NUDT21 HCT116 and shCPSF6-MDA cells to measure NEDD4 and NEDD4L mRNA isoform expression, which confirmed 3′ UTR shortening for both genes with CFIm loss (Fig. 4B-C). Next, we performed western blots to examine NEDD4 and NEDD4L protein expression in dTAG-NUDT21 HCT116 and shCPSF6-MDA cells under both CFIm intact and CFIm loss conditions. We found that CFIm loss increased NEDD4 and NEDD4L protein expression in both cell lines (Fig. 4D-E). To examine whether 3′ UTR shortening in NEDD4 and NEDD4L mRNAs contributes to the increase in their protein expression, we generated FL and short 3′ UTR reporters for both NEDD4 and NEDD4L and performed the dual fluorescence reporter assay as described earlier (Fig. 3D). For both NEDD4 and NEDD4L, the FL 3′ UTR strongly suppressed mCherry expression compared with the short 3′ UTR (mCherry/GFP: NEDD4-FL, 0.59; NEDD4-short, 0.95; NEDD4L-FL, 0.54; NEDD4L-short, 0.81) (Fig. 4F). Altogether, our results support that CFIm loss shortens the 3′ UTR of both NEDD4 and NEDD4L mRNAs and increases their protein expression.

**Figure 4.**
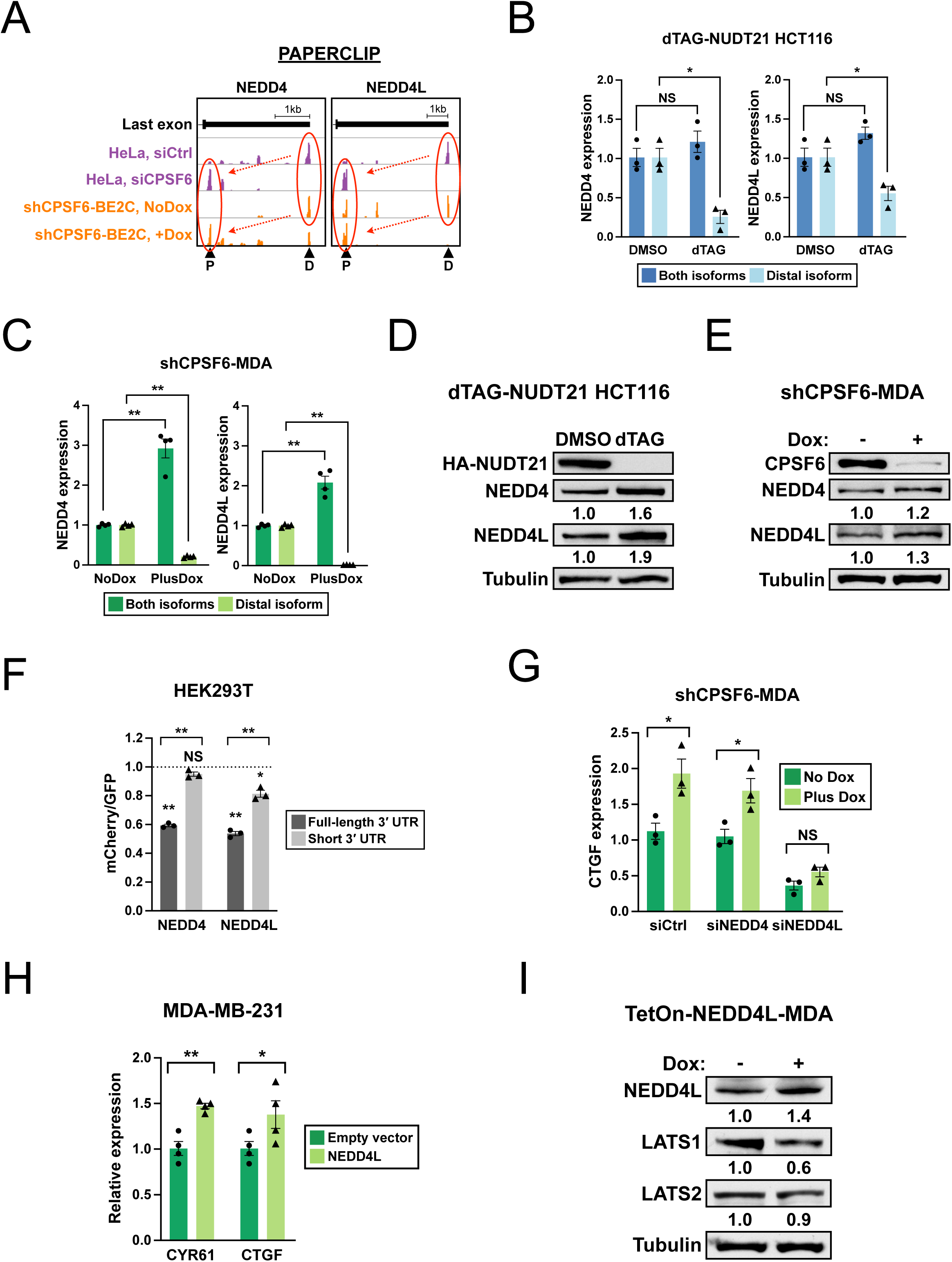
The identification of a CFIm-NEDD4L-LATS1 regulatory axis. (**A**) Results from our previously published PAPERCLIP experiments in HeLa cells and BE2C human neuroblastoma cells showing 3′ UTR shortening in NEDD4 and NEDD4L. Arrowheads: poly(A) sites identified by PAPERCLIP. P: Proximal. D: Distal. (**B**) Bar graphs showing expression of NEDD4 and NEDD4L mRNA isoforms measured by RT-qPCR in dTAG-NUDT21 HCT116 cells treated with DMSO or dTAG^V^-1 for 24 h from 3 independent experiments (n=3). (**C**) Bar graphs showing expression of NEDD4 and NEDD4L mRNA isoforms measured by RT-qPCR in shCPSF6-MDA cells treated with doxycycline (Plus Dox) or remained untreated (No Dox) for 72 h from 4 independent experiments (n=4). (**D∼E**) Western blots showing increased NEDD4 and NEDD4L expression in (**D**) dTAG-NUDT21 HCT116 cells treated with dTAG^V^-1 for 24 h, and (**E**) shCPSF6-MDA cells with doxycycline treatment for 72 h. Dox, doxycycline. Tubulin: loading control. (**F**) Bar graphs showing the mCherry/GFP ratio from NEDD4 and NEDD4L 3′ UTR reporters in 293T cells from 4 independent experiments (n=4). (**G**) Bar graphs showing CTGF expression measured by RT-qPCR in shCPSF6-MDA cells from 4 independent experiments (n=4). The cells were first transfected with control siRNAs (siCtrl), siRNAs targeting NEDD4 (siNEDD4), or siRNAs targeting NEDD4L (siNEDD4L) for 24 h and then were treated with doxycycline (Plus Dox) or remained untreated (No Dox) for 72 h before RNA collection. (**H**) Bar graphs showing CYR61 and CTGF expression measured by RT-qPCR in MDA-MB-231 cells transfected with an empty vector or an NEDD4L expression plasmid from 4 independent experiments (n=4). (**I**) Western blots showing decreased LATS1 expression in TetOn-NEDD4L-MDA cells with doxycycline treatment for 24 h. Dox, doxycycline. Tubulin: loading control. Error bars indicate SEM. Statistical significance is determined by one-tailed t-test (CTGF in panel H) and two-tailed t-test (all others). NS: not significant, *: p<0.05, **: p<0.01.

We next sought to examine the requirement of NEDD4 and NEDD4L in YAP/TAZ activation from CFIm loss in both dTAG-NUDT21 HCT116 and shCPSF6-MDA cells by simultaneously suppressing both genes using siRNAs (Fig. S3A-B). Double knockdown of NEDD4 and NEDD4L did not affect the increase in CTGF expression from CFIm loss in dTAG-NUDT21 HCT116 cells (Fig. S3C), but in shCPSF6-MDA cells, it indeed blunted the induction of CTGF from CFIm loss (Fig. S3D), indicating that NEDD4 and/or NEDD4L participated in YAP/TAZ activation from CFIm loss. To further examine which of the two genes are necessary for YAP/TAZ activation from CFIm loss, we next knocked down NEDD4 and NEDD4L individually by siRNAs in shCPSF6-MDA cells (Fig. S3E). We found that the induction of CTGF from CFIm loss was not affected by NEDD4 knockdown but was suppressed by NEDD4L knockdown (Fig. 4G), indicating that NEDD4L is necessary for YAP/TAZ activation from CFIm loss in shCPSF6-MDA cells.

We next transfected wildtype MDA-MB-231 cells with an NEDD4L-expressing plasmid, which induced CTGF and CYR61 expression (Fig. 4H), showing that NEDD4L overexpression alone is sufficient for YAP/TAZ activation. Lastly, because LATS1/2 are required for YAP/TAZ activation from CFIm loss in shCPSF6-MDA cells (Fig. 2G), we sought to explore a possible regulation between NEDD4L and LATS1/2. Therefore, we generated a stable MDA-MB-231 cell line that expresses NEDD4L under a doxycycline-inducible promoter (referred to as TetOn-NEDD4L-MDA cells hereafter), and we compared LATS1/2 protein expression in TetOn-NEDD4L-MDA cells between the presence and absence of doxycycline. We found that NEDD4L overexpression had modest effects on LATS2 but clearly lowered LATS1 expression (Fig. 4I), which suggests that NEDD4L can activate YAP/TAZ signaling through LATS1 suppression. In summary, we demonstrated that NEDD4L is both necessary and sufficient for YAP/TAZ activation from CFIm loss, and our results altogether support that CFIm suppresses YAP/TAZ signaling through NEDD4L and LATS1 in MDA-MB-231 cells and its derivatives.

### CFIm loss sensitizes cells to mRNA CPA inhibition

Because CFIm loss promotes PD-L1 expression and therapeutic resistance to several chemical inhibitors (Fig. 1F∼H), we next sought to find new therapeutic strategies for cancer with CFIm loss. JTE-607, which inhibits the endonuclease CPSF3 in the mRNA 3′-end processing complex, was recently shown to have anti-proliferation activities in some types of cancer such as acute myeloid leukemia (AML) and Ewing’s sarcoma^19^. Therefore, we treated dTAG-NUDT21 HCT116 cells with two different concentrations of JTE-607 in combination with either DMSO or dTAG^V^-1, and we compared cell proliferation after 48 h. Interestingly, CFIm loss enhanced the cytotoxic effects of JTE-607 at both concentrations (Fig. 5A), especially in the 10μM JTE-607 group, in which less than half the number of cells survived dTAG^V^-1 treatment compared to DMSO treatment (dTAG^V^-1: 37%, DMSO: 76%).

**Figure 5.**
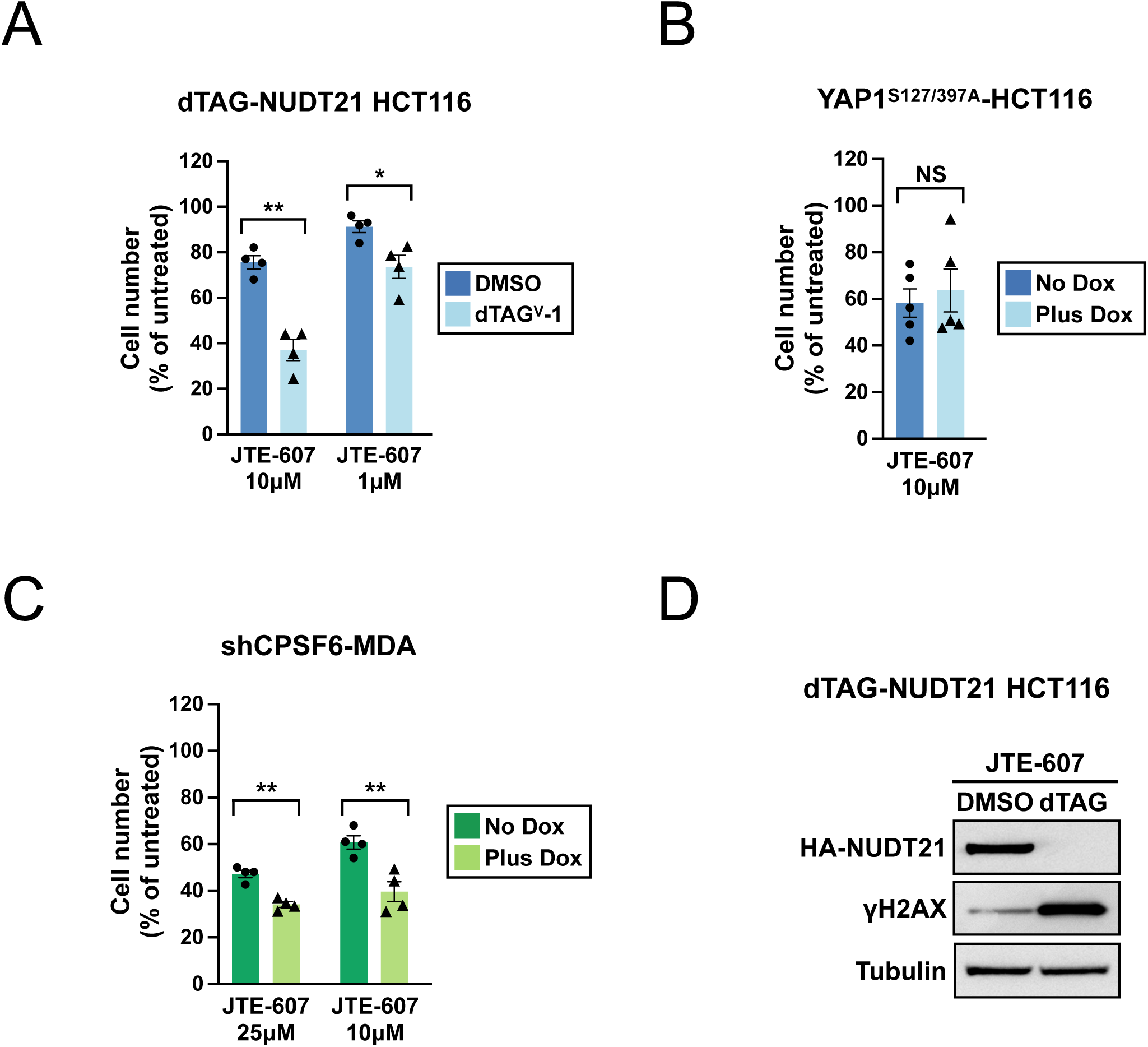
CFIm loss sensitizes cells to CPSF3 inhibition by exacerbating DNA damage. (**A**) Bar graphs showing the relative number of live dTAG-NUDT21 HCT116 cells treated with different concentrations of JTE-607 in combination with DMSO or dTAG^V^-1 for 48 h from 4 independent experiments (n=4). (**B**) Bar graphs showing the relative number of live YAP1^S127/397A^ HCT116 cells treated with 10μM JTE-607 alone (No Dox) or 10μM JTE-607 in combination with doxycycline (Plus Dox) for 48 h from 5 independent experiments (n=5). (**C**) Bar graphs showing the relative number of live shCPSF6-MDA cells treated with different concentrations of JTE-607 either alone or in combination with doxycycline for 72 h from 4 independent experiments (n=4). (**D**) Western blots showing increased γH2AX abundance in dTAG-NUDT21 HCT116 cells treated with 10μM JTE-607 and dTAG^V^-1 for 24 h. Tubulin: loading control. Error bars indicate SEM. Statistical significance is determined by two-tailed t-test. NS: not significant, *: p<0.05, **: p<0.01.

Next, we wished to determine whether YAP activation plays a role in the increased JTE-607 sensitivity from CFIm loss. To model YAP activation, we transduced HCT116 cells with a doxycycline-inducible YAP1^S127/397A^ transgene, which is resistant to Hippo pathway-mediated phosphorylation and inactivation^50^. Doxycycline treatment of YAP1^S127/397A^ HCT116 cells strongly activated CTGF expression (Fig. S4A) and increased Trametinib resistance (Fig. S4B), but it did not result in higher cytotoxicity with simultaneous JTE-607 treatment (Fig. 5B), suggesting that YAP activation does not contribute to the increased JTE-607 sensitivity from CFIm loss.

To address whether the increased vulnerability to JTE-607 from CFIm loss also extends to a different cell line, we next performed cell proliferation assay in shCPSF6-MDA cells. CFIm loss (Fig. S4C) again enhanced the cytotoxic effects of JTE-607 at both tested concentrations (Fig. 5C), with stronger effects in the 10μM JTE-607 group (Plus Dox: 40%, No Dox: 61%). Therefore, the increased vulnerability to JTE-607 from CFIm loss is not limited to HCT116 cells.

Because JTE-607 increased DNA damage in Ewing’s sarcoma cells^19^, we next examined whether stronger DNA damage contributes to the higher toxicity observed in dTAG-NUDT21 HCT116 cells receiving both JTE-607 and dTAG^V^-1 treatment (Fig. 5A). Western blot experiments showed a strong increase of gamma-H2AX, a widely used marker for DNA damage^51^, in dTAG^V^-1 treated cells when compared to DMSO treated cells (Fig. 5D). Altogether, our results support that the combination of CFIm loss and CPSF3 inhibition exacerbates DNA damage and causes cytotoxicity.

### CFIm loss and CPSF3 inhibition together induce BRCAness

We hypothesized that suppression of DNA damage repair (DDR) gene expression contributes to the stronger DNA damage in JTE-607 treated dTAG-NUDT21 HCT116 cells with CFIm loss. To comprehensively compare the expression of 276 DDR genes^52^ in JTE-607 treated dTAG-NUDT21 HCT116 cells between the DMSO (CFIm intact) and dTAG^V^-1 (CFIm loss) conditions, we performed RNA sequencing (RNA-seq) analysis (Fig. 6A). We found that two DDR pathways, the Fanconi anemia (FA) and homology-directed repair (HDR) pathways, were selectively affected by CFIm loss (Fig. 6B). Moreover, the gene expression changes for both pathways between the two conditions are not random. More than 80% of genes in the FA and HDR pathways decreased their expression in dTAG^V^-1 treated dTAG-NUDT21 HCT116 cells (Table S1).

**Figure 6.**
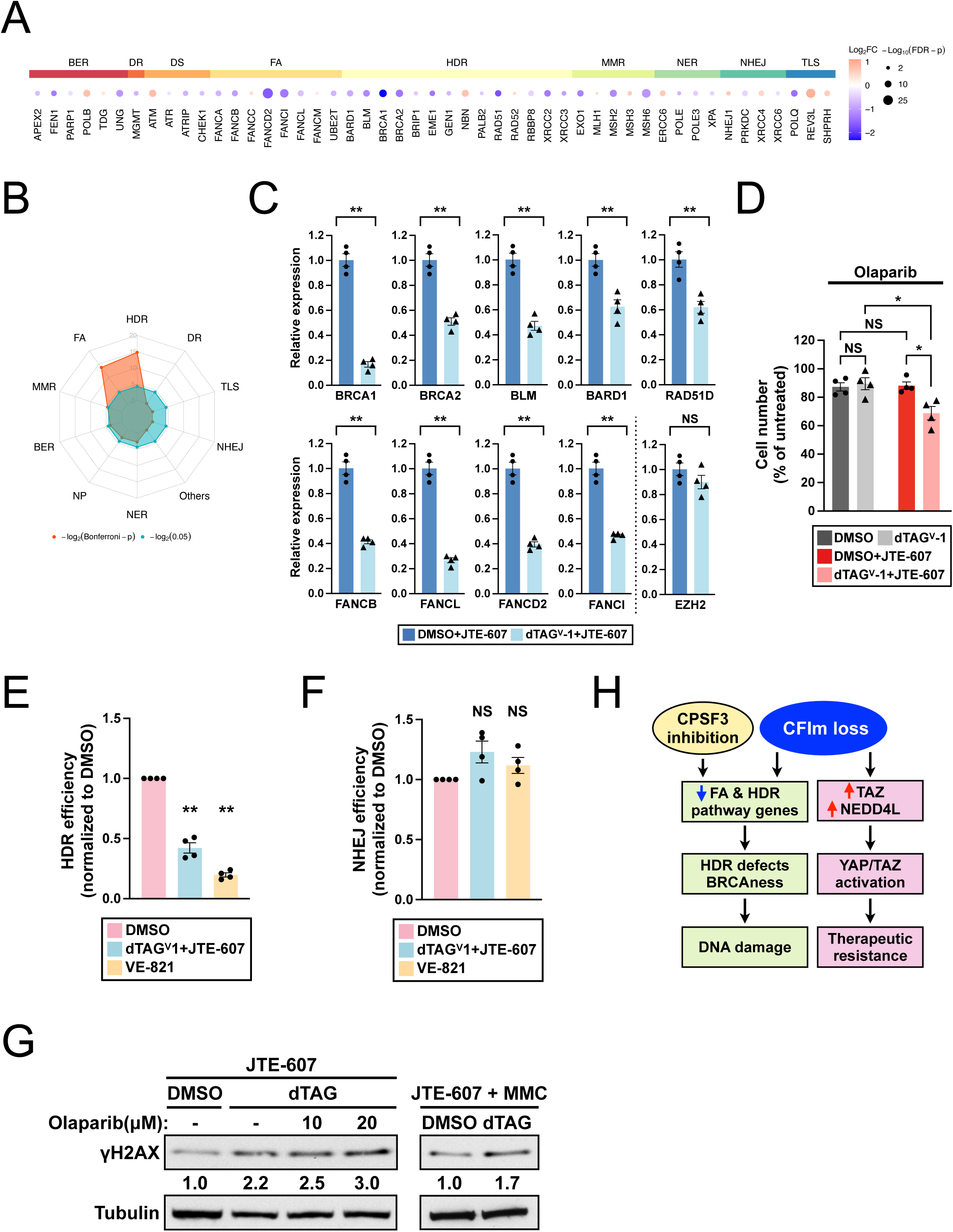
CFIm loss and CPSF3 inhibition together induce HDR defects and the BRCAness phenotype. (**A**) Heatmaps showing relative expression of core DDR genes from different DDR pathways between DMSO treated (CFIm intact) and dTAG^V^-1 treated (CFIm loss) dTAG-NUDT21 HCT116 cells in the presence of 10μM JTE-607 measured by RNA-seq from 4 independent experiments (n=4). (**B**) Radar plots showing relative enrichment of different DDR pathways in dTAG^V^-1 treated (CFIm loss) dTAG-NUDT21 HCT116 cells in the presence of 10μM JTE-607. (**C**) Bar graphs showing the expression of genes in the FA and HDR pathways measured by RT-qPCR from the same experiments in (A) (n=4). EZH2: negative control. (**D**) Bar graphs showing the relative number of live dTAG-NUDT21 HCT116 cells treated with 10μM Olaparib in combination with DMSO, dTAG^V^-1, DMSO plus 1μM JTE-607 or dTAG^V^-1 pllus 1μM JTE-607 for 24 h from 4 independent experiments (n=4). (**E and F**) Bar graphs showing the DNA repair efficiency of (E) the HDR pathway and (F) the NHEJ pathway measured using reporters in dTAG-NUDT21 HCT116 cells receiving either DMSO or dTAG^V^-1 plus 10μM JTE-607 from 4 independent experiments (n=4). VE-821, an ATR inhibitor that inhibits HDR, serves as a positive control in (E) and a negative control in (F). (**G)** Western blots showing elevated γH2AX abundance in dTAG-NUDT21 HCT116 cells treated with increasing concentrations of Olaparib (left panel) or with 100nM mitomycin C (MMC) (right panel) in the presence of 1μM JTE-607 and dTAG^V^-1 for 24 h. Tubulin: loading control. (**H**) Illustrations summarizing the cellular effects from CFIm loss. Error bars indicate SEM. Statistical significance is determined by two-tailed t-test. NS: not significant, *: p<0.05, **: p<0.01.

LOF mutations in HDR and FA pathway genes can cause HDR defects and the BRCAness phenotype^21,53^. Therefore, we hypothesized that CFIm loss and CPSF3 inhibition together induced the BRCAness phenotype in dTAG-NUDT21 HCT116 cells, and we performed a series of experiments to examine this hypothesis. We first performed RT-qPCR experiments to verify the suppression of 9 key genes in the FA and HDR pathways including *BRCA1/2* and several other BRCAness genes such as *RAD51D*^20,21^ and *FANCD2*^21^ (Fig. 6C) in dTAG-NUDT21 HCT116 cells receiving both dTAG^V^-1 and JTE-607. Notably, the expression of EZH2, which is not a DDR gene, remained at similar levels between DMSO and dTAG^V^-1 treated cells (Fig. 6C).

A key feature of the BRCAness phenotype is an increased sensitivity to PARP inhibitors. Thus, we next examined dTAG-NUDT21 HCT116 cells for their sensitivity to Olaparib, a PARP inhibitor. The Olaparib sensitivity was indistinguishable between DMSO treated and dTAG^V^-1 treated dTAG-NUDT21 HCT116 cells without JTE-607 (Fig. 6D, left). In contrast, Olaparib selectively suppressed proliferation of dTAG-NUDT21 HCT116 cells in the presence of both JTE-607 and dTAG^V^-1 (Fig. 6D, right). These results provide support that CFIm loss and CPSF3 inhibition together induced the BRCAness phenotype in dTAG-NUDT21 HCT116 cells.

Because the BRCAness phenotype is caused by HDR defects, we next performed DNA repair reporter assays^54,55^ using flow cytometry to examine whether the combination of JTE-607 and dTAG^V^-1 impairs HDR in dTAG-NUDT21 HCT116 cells. In these assays, successful DNA repair generates GFP expression, which reports the efficiency of the DNA repair pathway of interest. As expected, the combination of JTE-607 and dTAG^V^-1 treatment indeed strongly reduced HDR (Fig. 6E). In contrast, DNA repair by nonhomologous end-joining (NHEJ), a different DDR pathway that was not affected by JTE-607 and dTAG^V^-1 treatment in our RNA-seq analysis (Fig. 6B), remained intact (Fig. 6F).

Lastly, we performed western blot analysis to examine whether Olaparib is synergistic to JTE-607 and dTAG^V^-1 in inducing DNA damage in dTAG-NUDT21 HCT116 cells. Addition of Olaparib indeed elevated gamma-H2AX expression in dTAG-NUDT21 HCT116 cells receiving both JTE-607 and dTAG^V^-1 (Fig. 6G, left). Furthermore, mitomycin C (MMC), a DNA-crosslinking agent that BRCAness/HDR-deficient cancer is sensitive to^21^, also augmented DNA damage in dTAG^V^-1-treated dTAG-NUDT21 HCT116 cells in the presence of JTE-607 (Fig. 6G, right). Altogether, our results support that CFIm loss and CPSF3 inhibition together create a new vulnerability in cancer cells by impairing HDR and inducing the BRCAness phenotype (Fig. 6H).

## DISCUSSION

Activation of YAP/TAZ is common in human cancer and it often leads to therapeutic resistance and disease progression^2,4^. Therefore, understanding how cancer cells activate YAP/TAZ and their gene expression program is crucial to fight cancer therapy resistance. The Hippo pathway normally suppresses YAP/TAZ but the paucity of LOF mutations in the Hippo pathway components in cancer suggests the existence of alternative ways to activate YAP/TAZ. Our discovery that CFIm loss activates YAP/TAZ through the NEDD4L-LATS1 axis (Fig. 6H) has broad implications. First, it highlights the importance of post-transcriptional mechanisms in YAP/TAZ control, which is lesser known compared with the Hippo pathway. One example is the splicing factor ESRP2, which suppresses YAP activities by promoting the expression of *Yap1* and *Nf2* adult splicing isoforms in mouse liver^56^. We showed that CFIm suppresses YAP/TAZ activation by enforcing the use of full-length 3′ UTR in TAZ and NEDD4L (Fig. 3 and Fig. 4). Given the complexity of 3′ UTR regulation and the diverse types of cancer with YAP/TAZ activation^4,57^, future investigations might uncover additional mechanisms that are responsible for YAP/TAZ activation in cancer. Second, it shows that cancer cells activating YAP/TAZ through different mechanisms may have distinct responses to the same treatment. For example, we found that YAP mutation and CFIm loss both lead to YAP activation and Trametinib resistance in HCT116 cells (Fig. 1F and S4B). However, HCT116 cells with CFIm loss had higher resistance to YAP inhibitors but were more sensitive to JTE-607 (Fig. 1F and 5A). Therefore, separating cancer with high YAP/TAZ activities based on the YAP/TAZ activating mechanisms might be beneficial in revealing different treatment vulnerabilities.

Increasing evidence suggests that CPSF3 inhibition might be a viable treatment strategy for certain types of cancer such as AML, Ewing’s sarcoma, ovarian cancer and pancreatic ductal adenocarcinoma^17,19,58,59^. Conversely, CPSF3 inhibition seems to be less effective in many other types of cancer. For example, 92 cancer cell lines were tested for JTE-607 sensitivity based on cell viability in the original JTE-607 study^19^. Only 33 (36%) cell lines were sensitive to JTE-607 treatment (IC50 < 10μM), while 43 cell lines (47%) were classified as insensitive (IC50 > 20μM), including HCT-116 cells that we used in our current study. Our results thus demonstrate that it is possible to sensitize cancer cells to CPSF3 inhibition by inducing CFIm loss (Fig. 5A), which might widen the use of CPSF3 inhibition to more cancer types. Tian and colleagues recently reported that higher mRNA CPA activities resulted in stronger JTE-607 inhibition^17^. In their study, the cellular CPA activities were measured with a tandem PAS reporter, and an elevated CPA activity was designated as a preference to utilize the proximal PAS over the distal PAS^17^. Because CFIm loss results in widespread proximal APA shifts^10,12–14^, our results that CFIm loss increases JTE-607 sensitivity are therefore consistent with their proposed model of JTE-607 inhibition.

Inducing the BRCAness phenotype in HDR-proficient cancer cells has been shown to widen treatment options in breast and prostate cancer by increasing sensitivity to DNA damaging agents and PARP inihibitors^60,61^. Our study identified a new way to impair HDR and induce the BRCAness phenotype in cancer cells through the combination of CFIm loss and CPSF3 inhibition, and we provided multiple lines of evidence to support this finding (Fig. 6). Interestingly, a recent study showed that CPSF3 inhibition alone was sufficient to induce BRCAness phenotype through suppression of the HDR pathway in ovarian cancer^59^. Taken together, these results suggest that it is possible to induce the BRCAness phenotype in cancer cells through manipulating different components of the mRNA 3′ end processing complex, although the exact combination might vary in different types of cancer. Future studies to elucidate how different types of cells regulate their cellular CPA activities and their susceptibility to CPSF3 inhibition will provide important knowledge to improve cancer treatment.

Taken together, our study reveals a link between mRNA 3′ end processing and the YAP/TAZ transcription program. Despite the success in developing potent YAP/TAZ inhibitors, it remains challenging to treat YAP/TAZ activation in cancer because single treatment with YAP/TAZ inhibitors alone did not achieve sustained tumor suppression^35,36^. Our results suggest that the mRNA CPA inhibition might provide additional treatment benefits for cancer with CFIm loss. Besides cancer, the Hippo pathway and the YAP/TAZ transcription program also have important physiologic roles, especially in normal development and tissue regeneration^3^. Our results suggest that future studies on mRNA 3′ end processing and Hippo-YAP/TAZ in these contexts will provide important biological insights.

## Acknowledgements

We thank Amanda Clark for assistance in flow cytometry experiments. We thank the Health Sciences Sequencing Core at UPMC Children’s Hospital of Pittsburgh for high-throughput sequencing service, with special thanks to the assistant director, Will MacDonald. This work was supported by grants from the National Institutes of Health (R01NS113861 to H-W.H., R00CA207871 to H.U.O). Data analyses were supported by the University of Pittsburgh Center for Research Computing and the Extreme Science and Engineering Discovery Environment (XSEDE) supported by the National Science Foundation (OCI-1053575) via the Bridges2 system supported by the National Science Foundation (ACI-1445606) at the Pittsburgh Supercomputing Center.

## Author Contributions

Conceptualization, H.-W.H.; Methodology, A.N. and H.-W.H.; Formal analysis, J.H. and H.-W.H.; Investigation, M.G. and H.-W.H.; Writing—original draft preparation: H.-W.H.; Writing—review and editing: M.G., J.H., A.N., H.U.O., and H.-W.H.; Supervision: H.U.O. and H.-W.H.

## Declaration of interests

The authors declare no competing interests.

## STAR METHODS

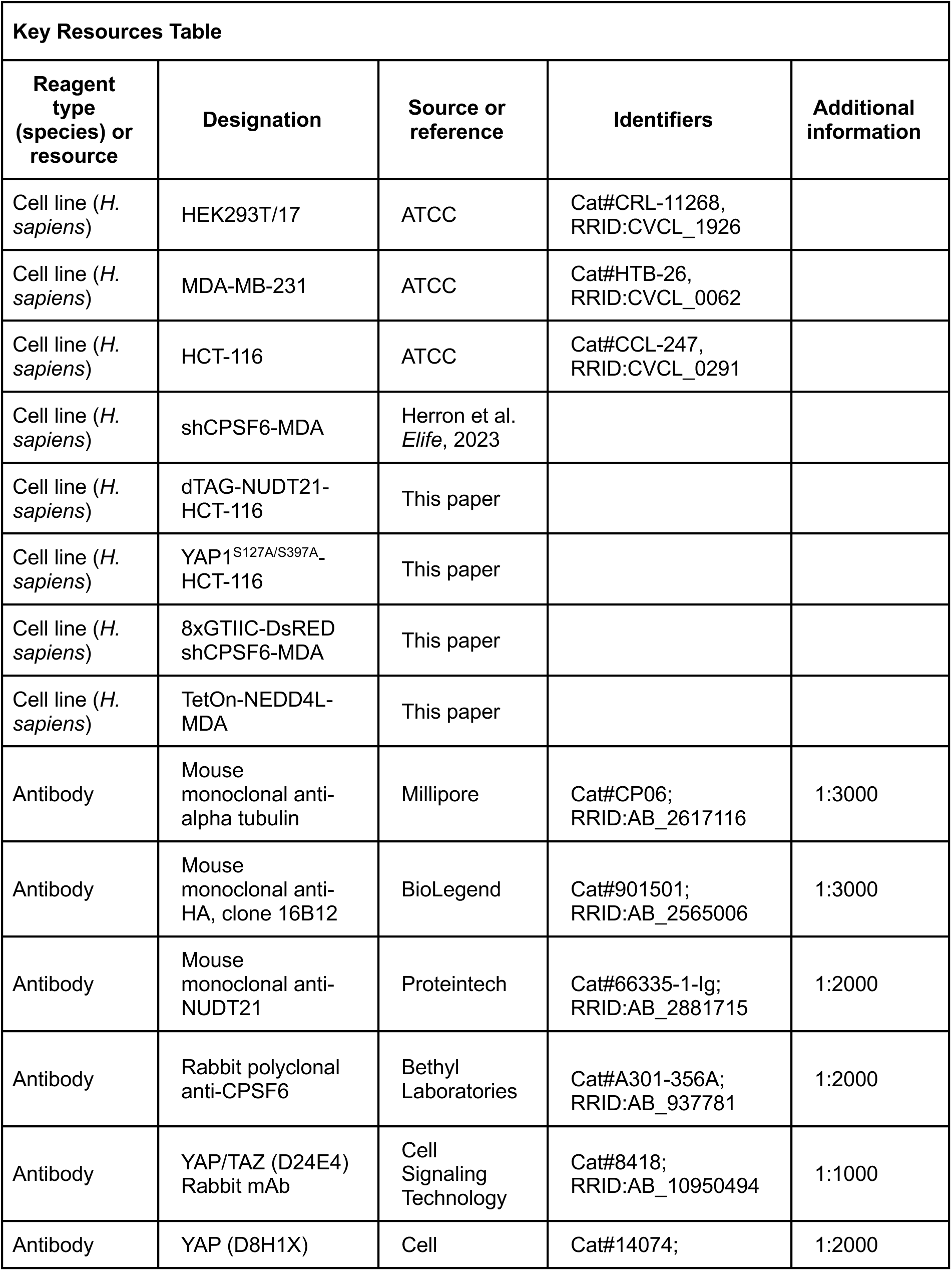
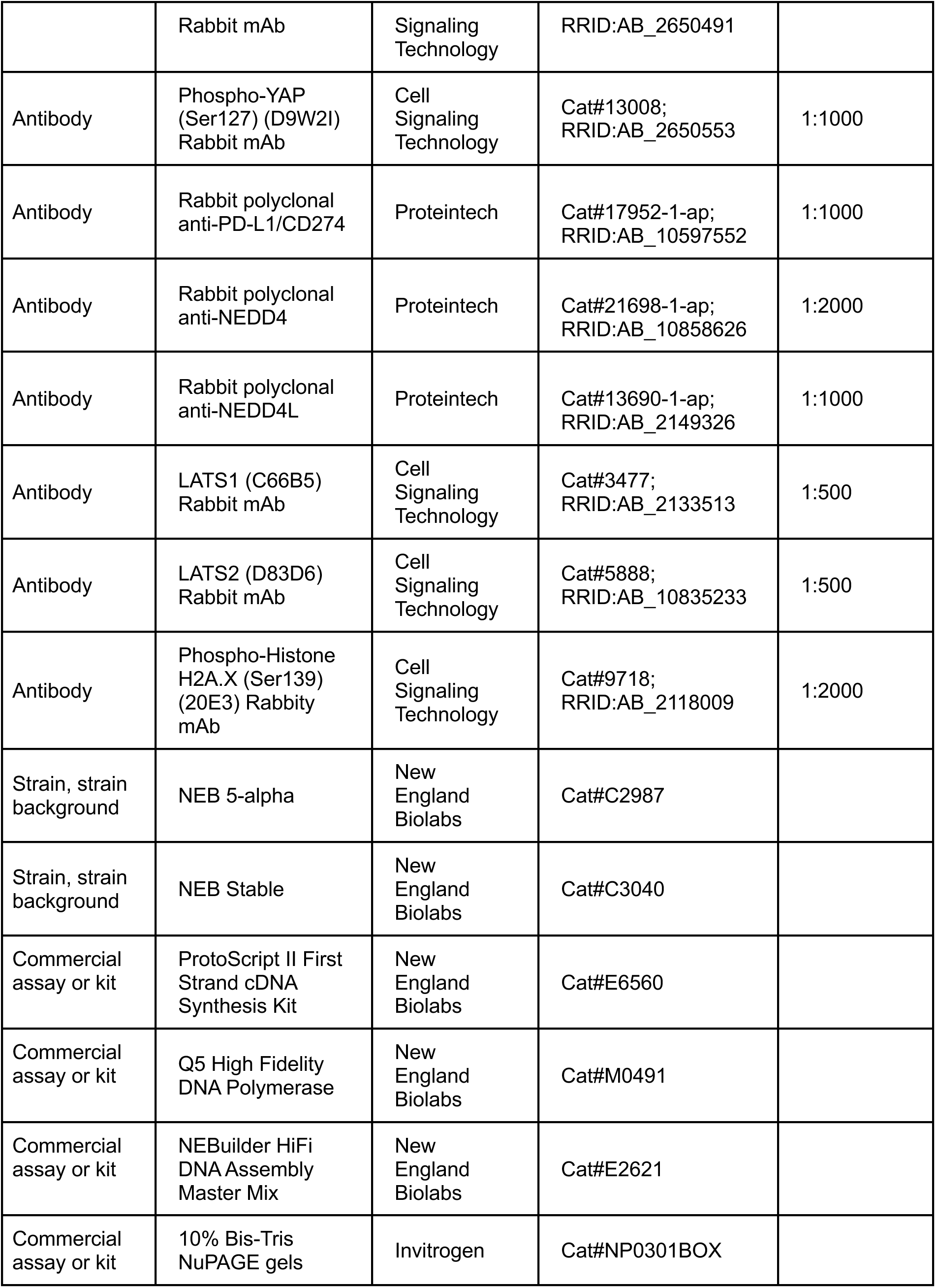
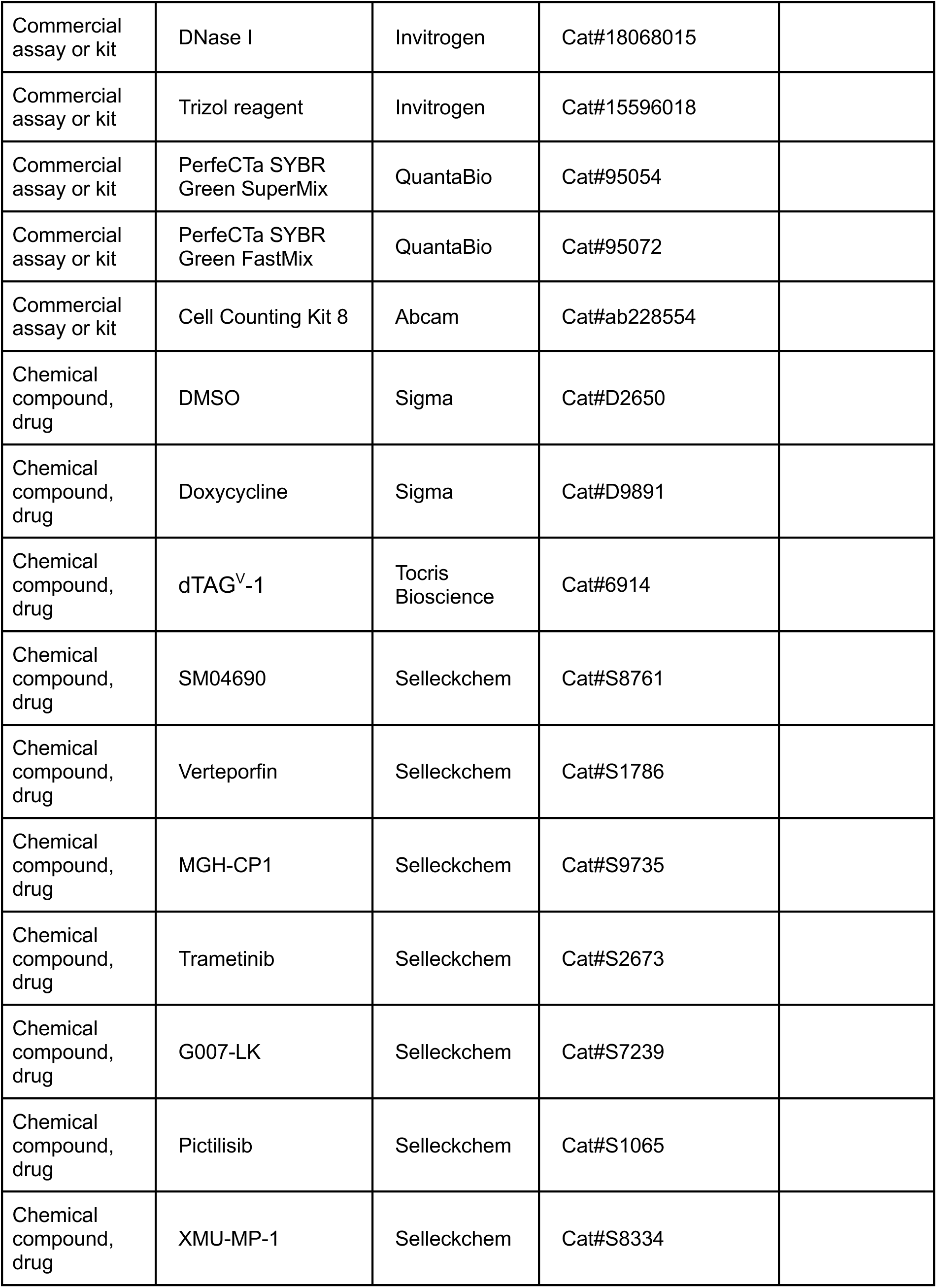
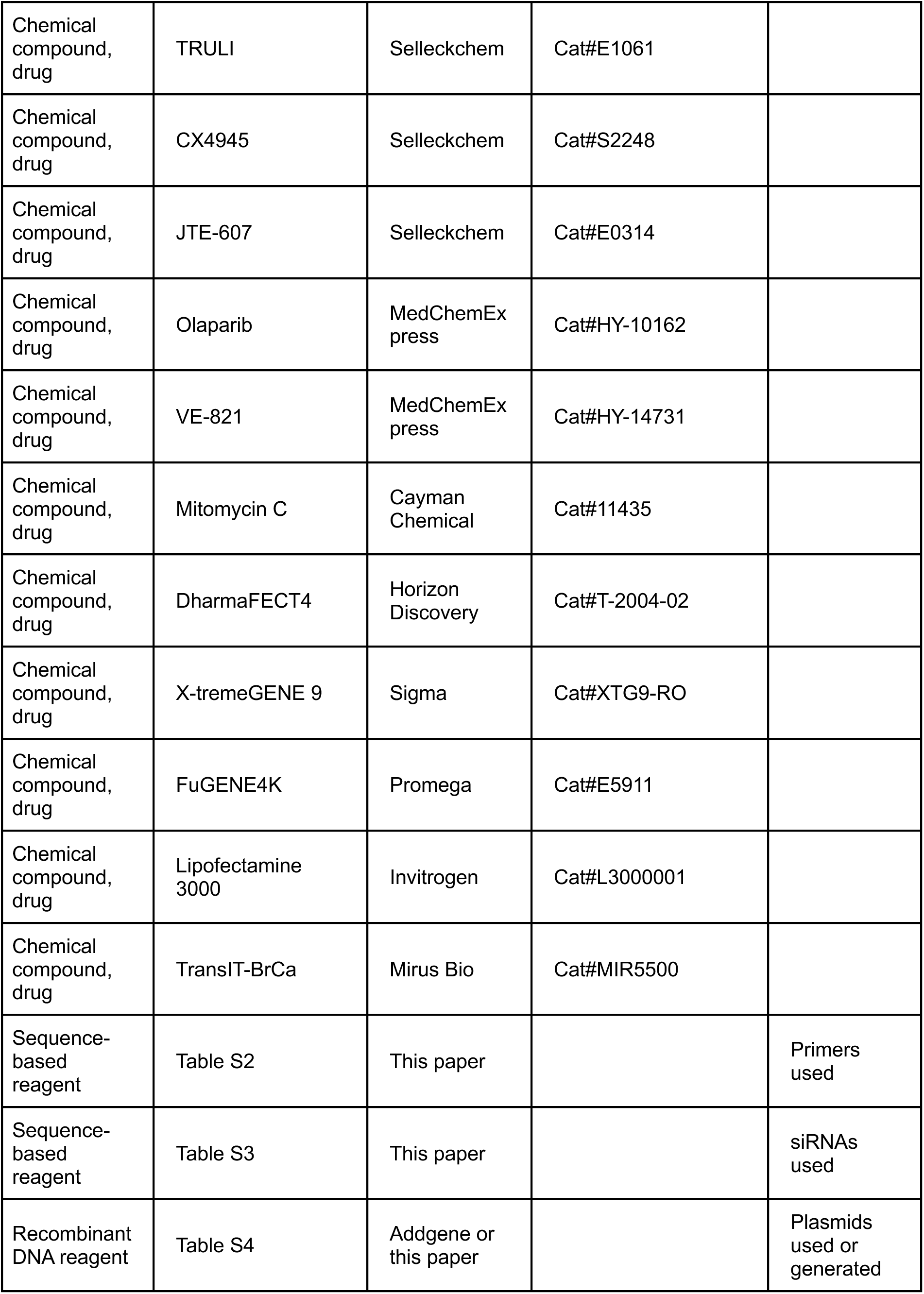
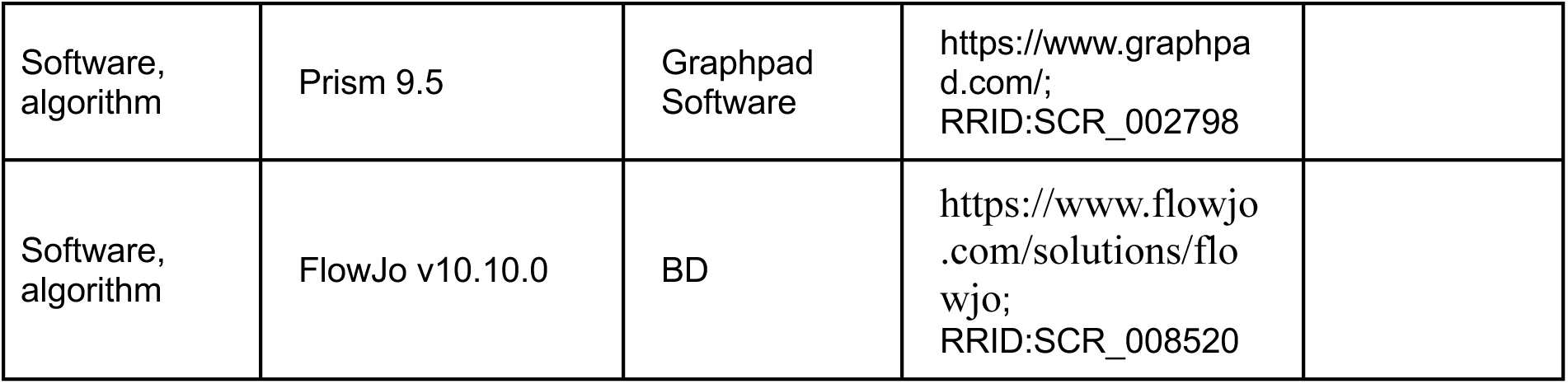

## RESOURCE AVAILABILITY

### Materials availability

Plasmids and cell lines generated for this study will be shared by the corresponding author upon request.

### Data availability

RNA-seq data have been deposited at GEO under the accession GSE306466.

## EXPERIMENTAL MODEL AND SUBJECT DETAILS

### Cell culture

HEK293T and MDA-MB-231 cells were grown in Dulbecco’s modified Eagle’s medium (DMEM). HCT116 cells were grown in McCoy’s 5A medium. All media were supplemented with 10% FBS and penicillin-streptomycin. Mycoplasma contamination in cell culture is screened with a commercial detection kit. HEK293T, MDA-MB-231, and HCT116 cells were obtained from ATCC with authentication and free of mycoplasma contamination. shCPSF6-MDA cells were previously generated in our lab^31^. 8xGTIIC-DsRED shCPSF6-MDA cells were generated by transducing shCPSF6-MDA cells with lentiviruses encoding an 8xGTIIC-DsRED YAP/TAZ reporter^32^ followed by hygromycin selection. TetOn-NEDD4L-MDA cells were generated by transducing rtTA3-expressing MDA-MB-231 cells with lentiviruses encoding doxycycline-inducible NEDD4L followed by puromycin selection. YAP1^S127/397A^ HCT116 cells were generated by transducing HCT116 cells with lentiviruses encoding doxycycline inducible YAP1^S127/397A^ followed by puromycin selection. dTAG-NUDT21 HCT116 cells were generated by CRISPR-mediated insertion of dTAG degron-containing cassettes into both *NUDT21* loci separately as previously described^26^. The transfected HCT116 cells were selected by Puromycin and Blasticidin, followed by single clone selection, DNA genotyping and Sanger sequencing to pick correctly inserted single clone cells. SiRNA transfection was performed using DharmaFECT4 (Horizon Discovery) with Silencer Select siRNAs (Invitrogen) at the final concentration of 25 nM following manufacturer’s instructions. All siRNAs used are listed in Table S2. Plasmid transfection was performed using X-tremeGENE9 (MilliporeSigma), FuGENE4K (Promega) and Lipofectamine 3000 (Invitrogen) following manufacturer’s instructions. Cell proliferation assays were performed in 96-well plates with Cell Counting Kit 8 (Abcam) following manufacturer’s instructions.

The following chemicals were used at the indicated concentrations: Doxycycline (Sigma): 1 μg/mL. dTAG^V^-1 (Tocris Bioscience): 500nM. Verteporfin (Selleckchem): 5μM. MGH-CP1 (Selleckchem): 10μM. Trametinib (Selleckchem): 10nM. G007-LK (Selleckchem): 10μM. Pictilisib (Selleckchem): 1μM. SM04690 (Selleckchem): 100nM. XMU-MP-1 (Selleckchem): 2μM. CX4945 (Selleckchem): 2μM. TRULI (Selleckchem): 10μM. Olaparib (MedChemExpress): 20μM. VE-821 (MedChemExpress0: 10μM. Mitomycin C (Cayman Chemical): 100nM.

## METHOD DETAILS

### RNA-seq

Library preparation of total RNA was carried out by using the Lexogen 3′ mRNA-Seq QuantSeq FWD V2 Kit according to manufacturer’s instructions. Library preparation, quality control, and sequencing were carried out Health Sciences Sequencing Core at UPMC Children’s Hospital of Pittsburgh. cDNA libraries were sequenced on an Illumina NextSeq machine (1 × 100 nt).

### Gene expression analysis

RNA-seq data were generated for four DMSO-treated (CFIm-intact) and four dTAGV-1–treated (CFIm-loss) dTAG-NUDT21 HCT116 cell samples. Raw FASTQ files underwent quality assessment with FastQC (https://www.bioinformatics.babraham.ac.uk/projects/fastqc/), evaluating per-base/tile sequence quality, adapter content, and duplication levels. Reads were preprocessed using fastp^62^ to trim auto-detected adapters and remove duplicates; sequences shorter than 25 bases after trimming were discarded. Clean reads were aligned to the Ensembl human reference genome (GRCh38, release 113) using STAR^63^. Gene-level counts were obtained with HTSeq^64^ by counting uniquely mapped reads overlapping annotated exonic regions.

Protein-coding genes with ≥10 counts in at least three samples were retained for downstream analysis. Count data were normalized using size factors estimated by DESeq2^65^, and a negative binomial generalized linear model was applied to identify differentially expressed genes (DEGs) between conditions. Genes with false discovery rate (FDR)-adjusted *p* < 0.05 were considered significant.

### Pathway enrichment analysis

Gene Set Enrichment Analysis (GSEA) was performed with clusterProfiler^66^ to evaluate enrichment of DNA damage response (DDR) pathways. Ten curated DDR gene sets were obtained from Knijnenburg et al.^52^. Gene sets with Bonferroni-adjusted *p* < 0.05 were considered significantly enriched.

### SDS-PAGE and western blots

30∼60 μg lysates from culture cells were separated on 10% Bis-Tris Novex NuPAGE gels (Invitrogen) and transferred to nitrocellulose membrane following standard procedures. The following antibodies are used for western blotting: mouse monoclonal anti-alpha tubulin (Millipore, CP06), mouse monoclonal anti-HA (clone 16B12, BioLegend, 901501), rabbit polyclonal anti-CPSF6 (Bethyl Labs, A301-356A), YAP/TAZ (D24E4) Rabbit mAb (Cell Signaling Technology, 8418), YAP (D8H1X) Rabbit mAb (Cell Signaling Technology, 14074), Phospho-YAP (Ser127) (D9W2I) Rabbit mAb (Cell Signaling Technology, 13008), Phospho-Histone H2A.X (Ser139) (20E3) Rabbity mAb (Cell Signaling Technology, 9718), mouse monoclonal anti-NUDT21 (Proteintech, 66335-1-Ig), rabbit polyclonal anti-PD-L1/CD274 (Proteintech, 17952-1-ap), rabbit polyclonal anti-NEDD4 (Proteintech, 21698-1-ap), rabbit polyclonal anti-NEDD4L (Proteintech, 13690-1-ap), LATS1 (C66B5) Rabbit mAb (Cell Signaling Technology, 3477), LATS2 (D83D6) Rabbit mAb (Cell Signaling Technology, 5888).

### Reverse transcription and quantitative PCR (RT-qPCR)

Quantitative PCR was performed using PerfeCTa SYBR Green SuperMix or PerfeCTa SYBR Green FastMix (QuantaBio) in triplicates. All primer sequences are listed in Table S2. For mRNA quantification, reverse transcription was performed using ProtoScript II First Strand cDNA Synthesis Kit (NEB) using d(T)_23_VN primer with DNase I (Invitrogen) digestion on 1 μg total RNA generated from Trizol (Invitrogen) extraction. The cycling parameters for qPCR were: 95°C for 10 min. followed by 40 cycles of 95°C for 15 sec., 58°C for 30 sec., 72°C for 20 sec.

Quantification was calculated using the ΔΔCt method with ACTB as the endogenous control. Normalization between replicate experiments was calculated by measuring ACTB expression with GAPDH as the endogenous control using the ΔΔCt method.

### Cloning and constructs

NEB HiFi DNA assembly following manufacturer’s instructions and standard cloning procedure (restriction digest, ligation, and transformation) was performed to generate desired constructs. All insert sequences were verified by Sanger sequencing. PITCh dTAG donor vectors for NUDT21 targeting were generated by insertion of custom guide RNA sequences using the following plasmids: pX330-BbsI-PITCh (Addgene, 127875), pCRIS-PITChv2-BSD-dTAG (BRD4) (Addgene, 91792), pCRIS-PITChv2-Puro-dTAG (BRD4) (Addgene, 91793). The following plasmids were obtained from Addgene and were used without modification: a lentiviral vector encoding doxycycline-inducible YAP1-S127/397A (Addgene, 213585), NEDD4 and NEDD4L expression plasmids (Addgene, 27002 & 27000). The YAP/TAZ reporter, pTRE-8xGTIIC-DsRED-DD-Hygro, was generated by replacing Neomycin in pTRE-8xGTIIC-DsRED-DD (Addgene, 115798) with Hygromycin. The dual fluorescence reporter for 3′ UTR assay (pcDNA5-FRT-GFP-mCherry-NoGW) was generated by removing the Gateway cloning fragment from pcDNA5-FRT-GFP-mCherry-3pGW (Addgene, 53965). The following 3′ UTR reporters were generated by insertion of 3′ UTR sequences amplified from human genomic DNA between the multiple cloning sites in pcDNA5-FRT-GFP-mCherry-NoGW: hTAZ-FL, hTAZ-S, hYAP-FL, hYAP-S, hNEDD4-FL, hNEDD4-S, hNEDD4L-FL, hNEDD4L-S.

### Dual fluorescence 3′ UTR reporter assay

For dual fluorescence 3′ UTR reporter assay, 293T cells were run on a FACSCanto II flow cytometer (BD) 24 h after transfection. The raw data were analyzed with FlowJo and then exported for calculation as previously described^67^. First, auto-fluorescence in both FITC and PE channels was calculated using untransfected cells, and the background is then defined as the mean plus 2 standard deviations of the auto-fluorescence signal. Next, for transfected cells, the measurements in both FITC and PE channels were subtracted with the background value, and cells with FITC and PE fluorescence levels below 0 after background subtraction were excluded from subsequent analyses. The data were next log-transformed and binned according to FITC levels, and the mean PE signal was calculated for each FITC bin. The average mCherry/GFP ratio was calculated using FITC bins between log2 to log4 with the empty vector set as 1.

### DNA repair reporter assays

For HDR reporter assay, dTAG-NUDT21 HCT116 cells were transfected with pDRGFP (Addgene, #26475) and pCBASceI (Addgene, #26477) together for 6 hours. The transfected cells then received different treatments for 18 hours before running through a FACSCanto II flow cytometer (BD) to determine the percentage of GFP-positive cells. For NHEJ reporter assay, dTAG-NUDT21 HCT116 cells were transfected with pimEJ5GFP (Addgene, #44026) and pCBASceI (Addgene, #26477) together for 24 hours. The transfected cells then received different treatments for 24 hours before running through a FACSCanto II flow cytometer (BD) to determine the percentage of GFP-positive cells. The raw data were analyzed with FlowJo (BD).

### QUANTIFICATION AND STATISTICAL ANALYSIS

Details of statistical tests are indicated below and in the Figure Legends. Statistical analyses were performed using R. For Figures 1C-G, 2B-G, 3B-C, 3E, 4B-C, 4F-G, 4H (CYR61), 5A-C, 6C-F, S1D-F, S2, S3, S4, statistical significance is determined by two-tailed Welch Two Sample t-test.

For Figures 4H (CTGF), statistical significance is determined by one-tailed Welch Two Sample t-test.

For all figures: *: p < 0.05, **: p < 0.01.

## SUPPLEMENTARY FIGURE LEGENDS

**Figure S1.**
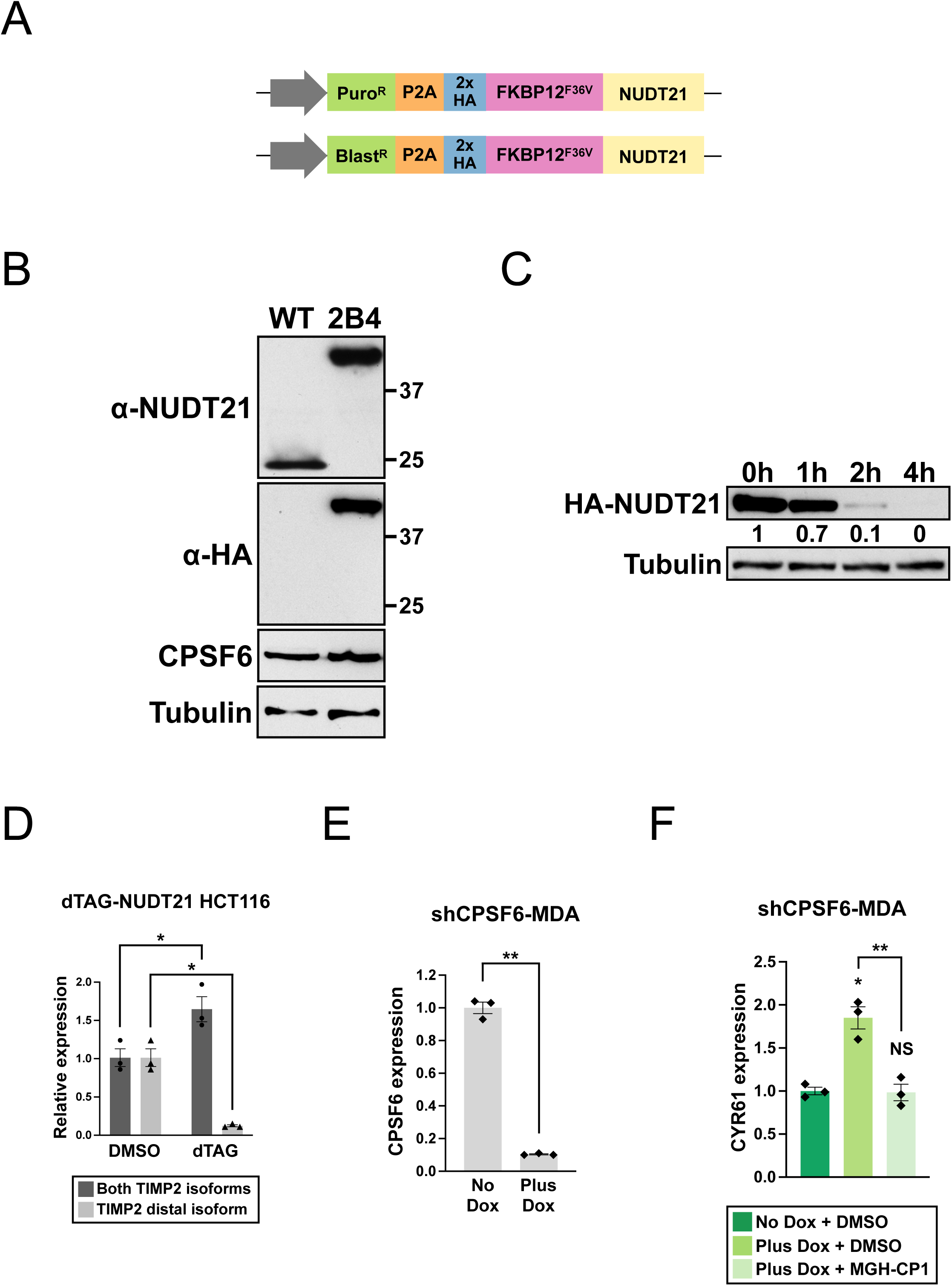
Characterization of dTAG-NUDT21 HCT116 cells, related to Figure 1. (**A**) An illustration showing the *NUDT21* loci in dTAG-NUDT21 HCT116 cells after CRISPR-mediated insertion of the dTAG degron-containing cassettes. Arrow: the endogenous *NUDT21* promoter. Puro^R^: puromycin resistance gene. Blast^R^: Blasticidin resistance gene. (**B**) Western blots showing the expression of N-terminal tagged NUDT21 and endogenous CPSF6 expression in dTAG-NUDT21 HCT116 cells, which are derived from a single cell clone (2B4). α-NUDT21: NUDT21 antibody; α-HA: HA antibody. WT: wildtype HCT116 cells. (**C**) Western blots showing a complete loss of NUDT21 in 4 h dTAG^V^-1 treated dTAG-NUDT21 HCT116 cells. Tubulin: loading control. (**D**) Bar graphs showing TIMP2 isoform expression measured by RT-qPCR in dTAG-NUDT21 HCT116 cells treated with DMSO or dTAG^V^-1 for 24 h from 3 independent experiments (n=3). (**E**) Bar graphs showing CPSF6 expression measured by RT-qPCR in shCPSF6-MDA cells treated with doxycycline (Plus Dox) or remained untreated (No Dox) for 72 h from the same 3 independent experiments shown in Fig. 1D (n=3). (**F**) Bar graphs showing CYR61 expression measured by RT-qPCR in shCPSF6-MDA cells receiving different treatments for 72 h from the same 3 independent experiments shown in Fig. 1G (n=3). dTAG^V^-1: 500nM. Doxycycline: 1 μg/mL. MGH-CP1: 10μM. Error bars indicate SEM. Statistical significance is determined by two-tailed t-test. NS: not significant, *: p<0.05, **: p<0.01.

**Figure S2.**
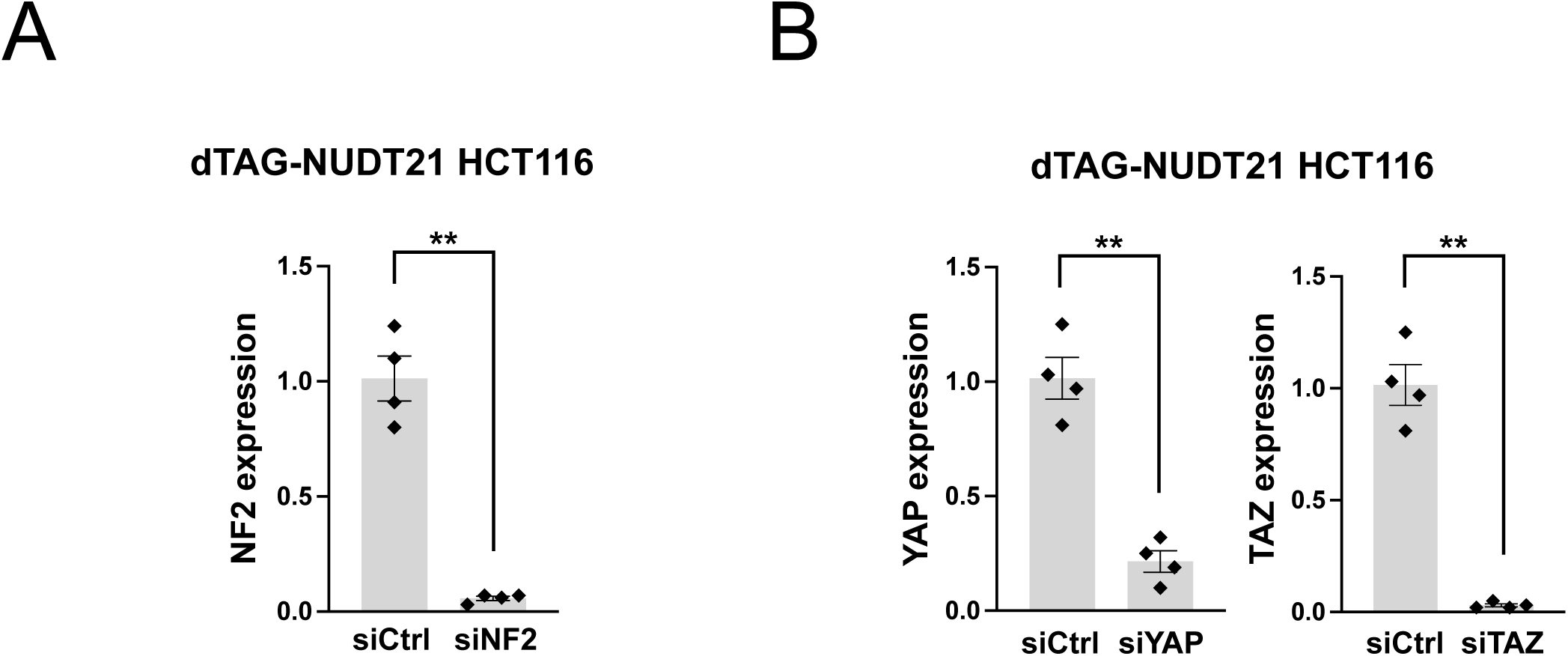
siRNA knockdown efficiency in dTAG-NUDT21 HCT116 cells, related to Figure 2. (**A**) Bar graphs showing NF2 expression measured by RT-qPCR in dTAG-NUDT21 HCT116 cells from the same 4 independent experiments shown in the right panel of Fig. 2C (n=4). Cells were first transfected with control siRNAs (siCtrl) or siRNAs targeting NF2 (siNF2) for 48 h and then were treated with DMSO or dTAG^V^-1 for 24 h before RNA collection. (**B**) Bar graphs showing YAP and TAZ expression measured by RT-qPCR in dTAG-NUDT21 HCT116 cells from the same 4 independent experiments shown in Fig. 2E (n=4). Cells were first transfected with control siRNAs (siCtrl), siRNAs targeting YAP (siYAP), or siRNAs targeting TAZ (siTAZ) for 48 h and then were treated with DMSO or dTAG^V^-1 for 24 h before RNA collection. dTAG^V^-1: 500nM. Doxycycline: 1 μg/mL. Error bars indicate SEM. Statistical significance is determined by two-tailed t-test. **: p<0.01.

**Figure S3.**
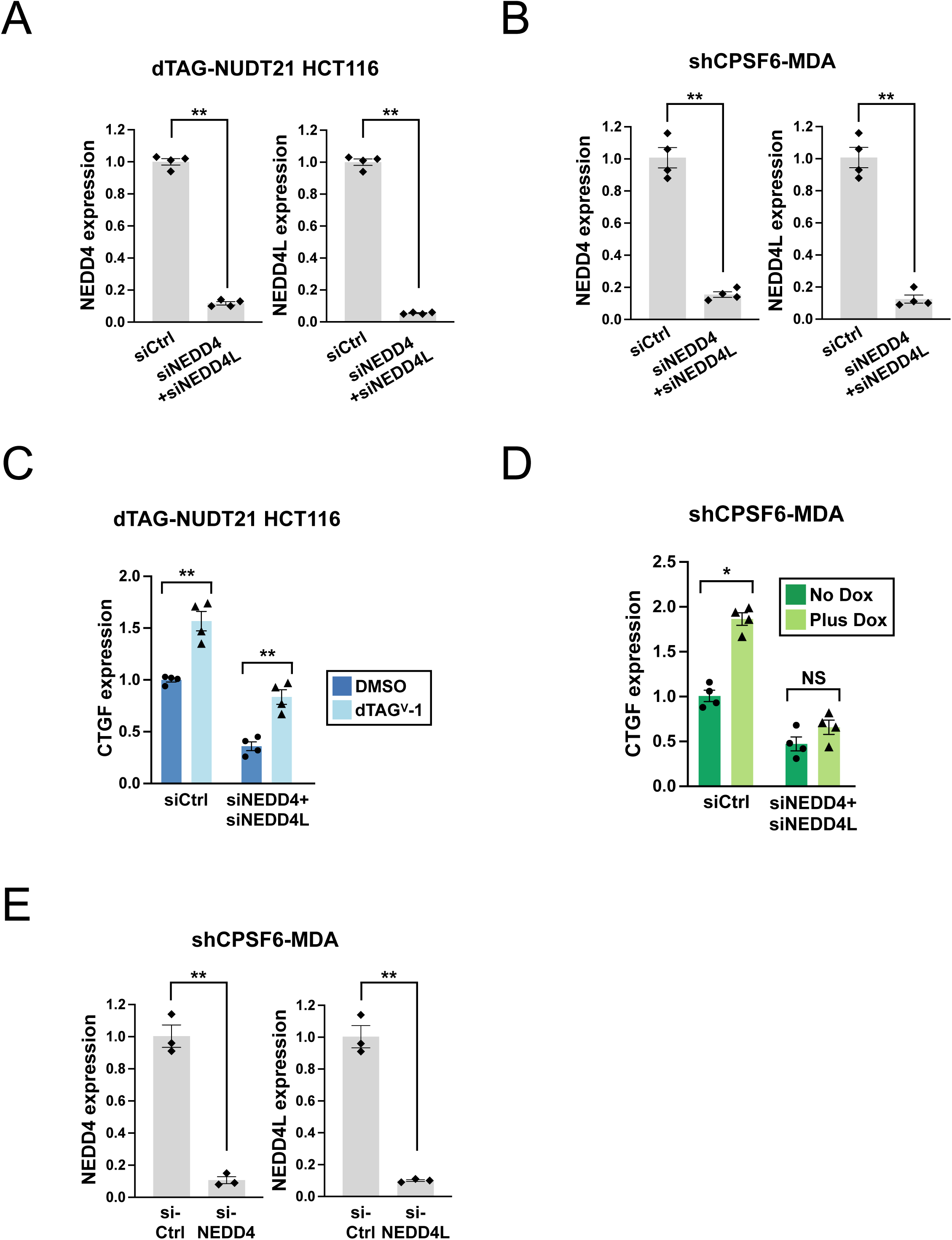
Examining YAP/TAZ activation in shCPSF6-MDA and dTAG-NUDT21 HCT116 cells with NEDD4 and NEDD4L double knockdown, related to Figure 4. (**A**) Bar graphs showing NEDD4 and NEDD4L expression measured by RT-qPCR in dTAG-NUDT21 HCT116 cells from 4 independent experiments (n=4). The cells were first transfected with control siRNAs (siCtrl) or siRNAs targeting both NEDD4 (siNEDD4) and NEDD4L (siNEDD4L) for 48 h and then were treated with DMSO or dTAG^V^-1 for 24 h before RNA collection from 4 independent experiments (n=4). (**B**) Bar graphs showing NEDD4 and NEDD4L expression measured by RT-qPCR in shCPSF6-MDA cells from 4 independent experiments (n=4). The cells were first transfected with control siRNAs (siCtrl) or siRNAs targeting both NEDD4 (siNEDD4) and NEDD4L (siNEDD4L) for 24 h and then were treated with doxycycline (Plus Dox) or remained untreated (No Dox) for 72 h before RNA collection. (**C**) Bar graphs showing CTGF expression measured by RT-qPCR in dTAG-NUDT21 HCT116 cells from the same 4 independent experiments shown in panel A (n=4). (**D**) Bar graphs showing CTGF expression measured by RT-qPCR in shCPSF6-MDA cells from the same 4 independent experiments shown in panel B (n=4). (**E**) Bar graphs showing NEDD4 or NEDD4L expression measured by RT-qPCR in shCPSF6-MDA cells from the same 3 independent experiments shown in Fig. 4G (n=3). Error bars indicate SEM. Statistical significance is determined by two-tailed t-test. NS: not significant, *: p<0.05, **: p<0.01.

**Figure S4.**
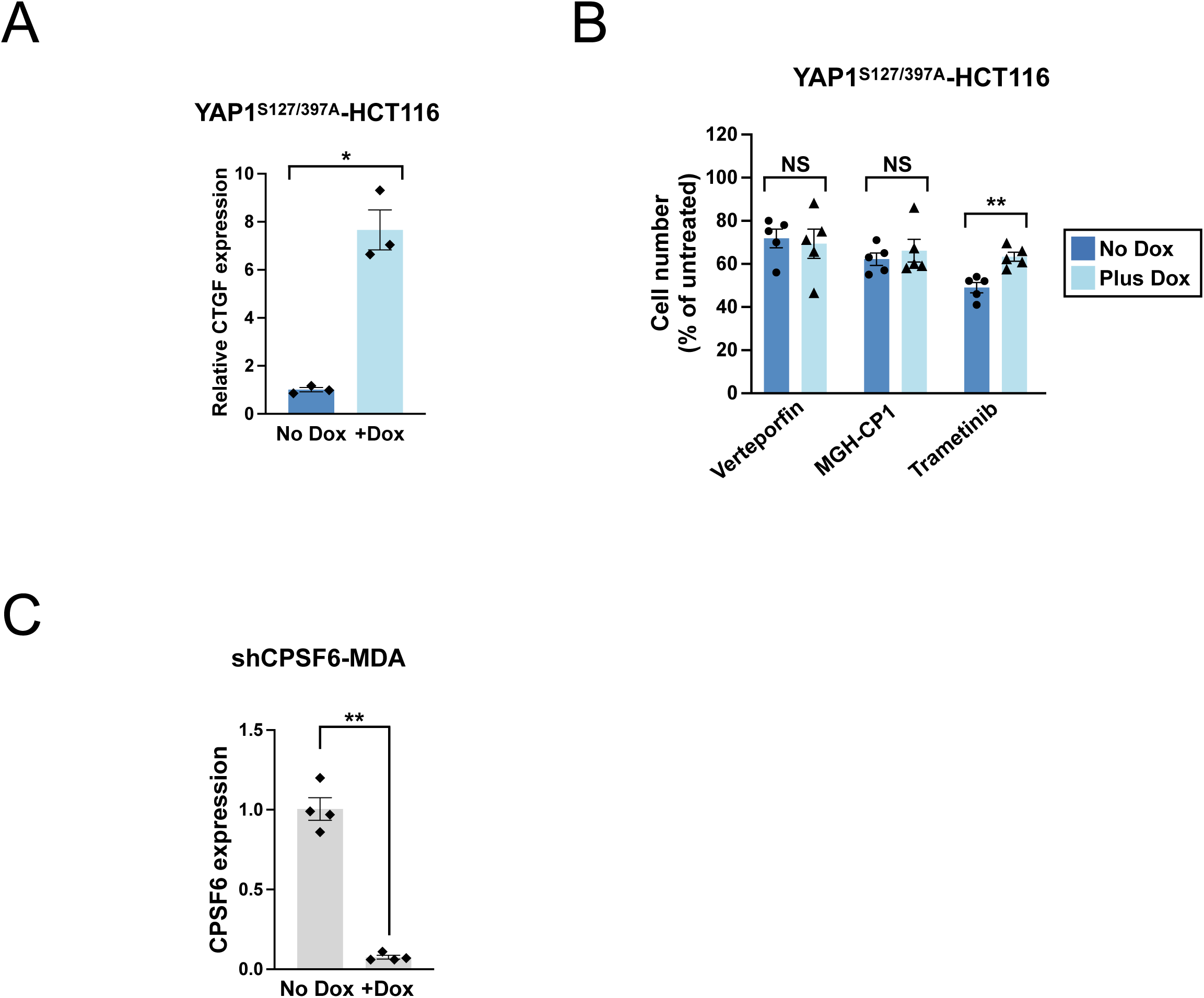
Doxycycline treatment of YAP1^S127/397A^ HCT116 cells activates YAP signaling and promotes Trametinib resistance, related to Figure 5. (**A**) Bar graphs showing CTGF expression measured by RT-qPCR in YAP1^S127/397A^ HCT116 cells treated with treated with doxycycline (+Dox) or remained untreated (No Dox) for 24 h from 3 independent experiments (n=3). (**B**) Bar graphs showing the relative number of live YAP1^S127/397A^ HCT116 cells treated with different chemical inhibitors, either alone (No Dox) or in combination with doxycycline (Plus Dox) for 48 h from 5 independent experiments (n=5). (**C**) Bar graphs showing CPSF6 expression measured by RT-qPCR in shCPSF6-MDA cells treated with doxycycline (+Dox) or remained untreated (No Dox) for 72 h from the same 3 independent experiments shown in Fig. 5G (n=4). Doxycycline: 1 μg/mL. Verteporfin: 5μM. MGH-CP1: 10μM. Trametinib: 10nM. Error bars indicate SEM. Statistical significance is determined by two-tailed t-test. NS: not significant, *: p<0.05, **: p<0.01.

## LIST OF NON-PDF SUPPLEMENTAL ITEMS

**Table S1.** DDR gene expression changes from RNA-seq profiling, related to Figure 6.

**Table S2.** List of primers, related to Methods.

**Table S3.** List of siRNAs, related to Methods.

**Table S4.** List of plasmids, related to Methods.

## Notes

### Competing Interest Statement

The authors have declared no competing interest.

